# Noise Optimization of Basic Signal Component Extraction for Cryogenic and On-Scalp Magnetoencephalography (MEG)

**DOI:** 10.64898/2026.07.21.739883

**Authors:** Alexandria McPherson, Sepp Sanchirico, Albert Xu, Eric Larson, Milena Kaestner, William Turner, Laura Gwilliams, Samu Taulu

## Abstract

Magnetoencephalography (MEG) measures human neural activity non-invasively with spatio-temporal precision, and has been foundational in enabling impactful discoveries in cognitive neuroscience. New on-scalp MEG sensor technologies, such as OPM-MEG, offer the opportunity to capture more information about the neuronal magnetic fields with higher sensitivity to more complex, higher order spatial components, leading to improved source localization. The accuracy of MEG and OPM-MEG source localization relies on data preprocessing techniques to isolate the neuronal fields from other magnetic and biomagnetic sources through signal space separation, rejection, and suppression methods. Current preprocessing methods risk rejecting brain signals of interest, or can spread sensor noise artifacts unknowingly. Here we propose a novel preprocessing method for MEG, and test the extent to which it overcomes limitations of prior methods. Specifically, we derive, apply, and assess a novel signal space separation (SSS) method with Foster’s inverse, a weighted matrix inversion protocol that can utilize information about the MEG sensor noise and artifacts to reconstruct neuronal activity. With simulations, phantom head cryogenic MEG recordings, and subject recordings with two OPM-MEG systems, we show that Foster’s inverse with SSS offers a more robust and stable reconstruction of the neuronal magnetic fields, especially in the face of sensor noise and artifacts. As the field of cognitive neuroscience continues to embrace MEG and OPM-MEG, Foster’s inverse with SSS offers a robust and powerful data preprocessing technique for reducing noise and improving source localization of the underlying neuronal currents.

## 1 Introduction

### 1.1 Motivation

Advancements of non-invasive brain imaging techniques have allowed cognitive neuroscience to study the function, development, and structure of the healthy human brain, leading to impactful discoveries in human neuroscience [15], [3], [6], [20], [28], [30]. Cryogenic magnetoencephalography (MEG) and optically pumped magnetoencephalography (OPM-MEG) are non-invasive imaging techniques that provide great temporal and good spatial precision of brain activity. These systems measure the weak magnetic fields produced by neuronal activity [16], from sensors placed on or near the scalp. The distance between the sensors and the neural populations of interest introduces two primary challenges for MEG. The first is to recover robust signal strength, because the amplitude of the signal decreases with distance from the electromagnetic source. The second is to accurately localize where brain activity is coming from, which needs to be inferred from the spatial and temporal patterns of activity recorded at the scalp. Separating signal from noise, and retaining fidelity of spatial information, is therefore a critical and determining factor for constraining the research questions that can be addressed, and conclusions that can be drawn, from MEG data.

Several algorithms have been developed to improve signal-to-noise ratio (SNR) in time-resolved neural signals such as MEG. Some of these preprocessing methods rely on spatial projections or frequency filters to remove noise [41]; others serve to suppress sensor noise specifically [5], [23]; and others utilize knowledge of expected magnetic field properties to separate neuronal fields from unwanted magnetic sources [27], [34], [35], [18], [39], [38], [37]. While these methods serve to improve SNR when certain conditions are met, they also have limitations. For example, spatial projections and frequency filtering run the risk of removing signals of interest during the noise removal process. Furthermore, when the sensor data contain a large noise source, this can lead to corruption of the signal with unanticipated bleed-over from the noise. Finally, some signal separation methods require high numbers of MEG sensors to get enough samples of the spatiotemportal patterns of the signal, which can lead to increased instability for systems with a low channel count.

Here we describe, apply, and assess an algorithm for signal component extraction for MEG data. We show that in theory, in simulation, and in empirical data, our approach overcomes many of the challenges faced by extant methods. The success of our approach can be attributed to a novel signal space separation algorithm that weights the solution for the underlying neuronal activity with information describing the sensor noise and artifacts. This enables a stable solution that is robust against large noise sources and depends on few assumptions. It also boosts source localization accuracy by being sensitive to the higher spatial components in the signal, when tested on a phantom dataset. In what follows, we outline the method and its theory, and step through both simulation results under different noise conditions, and empirical results when applied to two OPM-MEG datasets of auditory evoked responses. The method and results are applicable to all off- and on-scalp MEG systems, but we will focus on cryogenic MEG and OPM-MEG here.

### 1.2 Magnetometers for Neuroimaging

Magnetic fields generated by the brain typically range from 10 fT to 1 pT in strength, and 1-100 Hz in frequency [16]. These fields are mainly produced by the activity of pyramidal cortical neurons aligned perpendicular to the gray/white matter boundary, where activity of 10,000 to 50,000 neurons is modeled as a current dipole [16], [30], [33]. Based on the magnetic field distribution measured by sensors arranged around the head, the underlying neural current can be reconstructed.

Traditional MEG sensors are made of an array of superconducting quantum interference devices (SQUIDs) that are kept in the superconducting state by cooling them with liquid helium in a Dewar elevated above the patient’s scalp [16], with approximately 18 mm between the room-temperature surface of the MEG helmet and the SQUID sensors themselves. Due to the spatial degradation of magnetic fields as a function of distance and the presence of numerous external noise sources, it is difficult to accurately measure these small brain signals. Additionally, noise from external sources, movement of the patient, sensor noise and artifacts, and any remnant magnetic fields around the sensors can cause signal-like artifacts that are difficult to distinguish from brain activity itself [37].

Newer on-scalp sensors, such as optically pumped magnetometers (OPM), have recently begun to be implemented in MEG systems [1], [32], [40] [19], [4]. OPM sensors exploit the quantum-mechanical properties of alkali metal gas, typically Rubidium, within the sensors [1]. Energy is pumped in from a modulating laser to bring the gas atoms into the same energy level such that they are at their most sensitive to small changes in magnetic fields [40]. In the presence of neuronal fields, the spin momentum and angular momentum of the atoms are disrupted, allowing them to once again reabsorb light and energy from the pumping laser to realign the state of the atoms. Under ideal conditions, called the spin-exchange relaxation-free (SERF) conditions, the amplitude of laser light measured through the sensor is linearly related to the magnetic field along the laser axis, and gives OPMs the ability to localize sources within 2-3 mm [4]. Specifically, SERF requires a high concentration of gas within the sensor cell to help maintain a consistent average angular momentum across all particles, and requires the background magnetic field to be sufficiently low (*<* 5 nT) to maintain the desired spin momentum of the valence electron in each atom [1], [4], [40].

Because OPMs operate at room temperature and can be worn on the scalp, they offer a three- to five-fold improvement in sensitivity to neuronal magnetic fields, and are able to capture higher order spatial components of the magnetic field, which decay more rapidly over the space between neuronal sources and sensors [21], [42], [42],[40], [12]. In the context of cognitive neuroscience, these advancements directly relate to increased SNR and increased localization accuracy [4]. Furthermore, the ability of OPM sensors to operate without being submerged in liquid helium gives participants freedom to move with the MEG array fixed to their head, enabling ambulatory neuroscience experiments [42]. However, due to the requirement that OPM sensors operate under 5nT background magnetic fields to maintain SERF conditions, OPM-MEG systems experience sensitivity to low-frequency fields that cryogenic MEG systems are more robust to [4], [40]. Additionally, the potential for higher crosstalk between sensors, localization and calibration errors from variable sensor positions, and patient movement with the sensor helmet can contribute to sensor noise or drift artifacts in OPM-MEG recordings [4], [40]. Often in MEG and EEG, these low frequency fields can be high-pass filtered out, but raising these filters above 0.5Hz can eliminate low-frequency brain activity of interest [8]. By mathematically accounting for the sensor noise, artifacts, and external magnetic fields present in both OPM-MEG and cryogenic MEG recordings, the algorithm we introduce here - Foster’s inverse with SSS - accurately reconstructs the underlying neuronal activity with reduced impacts from sensor noise.

### 1.3 Related Signal Processing Approaches

The foundational methods for isolating neuronal magnetic fields are based on the physics of electromagnetism. Brain signals can be separated from external interference signals using, for example, the signal space separation (SSS) method (also known as Maxwell filtering). SSS is a foundational method for the discretization of magnetic fields, and works by modelling the quasi-static magnetic fields as a series of vector spherical harmonics (VSH), with separate terms for signal contributions that originate inside the brain (“interior basis”) and outside the brain (“exterior basis”) [34]. SSS is based only on the geometric configuration of the MEG sensor helmet in relation to the neuronal sources [36]. The present magnetic fields are continuous in space and time, but are discretely sampled by the MEG sensors. As such, they can be represented as a sum of increasingly complex spatial topographies. Figure 1 provides an example of the increased complexities with increased discrete orders of *L*, the precise mathematical definition of which will be unpacked in Section 2. How many discrete orders of *L* to use - i.e., where to truncate the discrete expansion of the magnetic field - depends on factors such as the number of MEG sensors, and the distance between sensors and scalp. These parameters, of course, are quite different between cryogenic-MEG and OPM-MEG systems [36].

**Figure 1.**
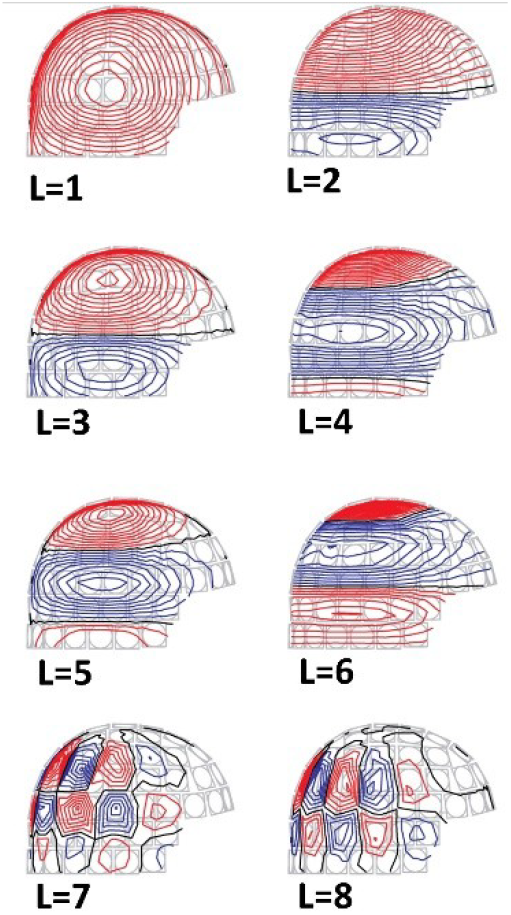
Low to high spatial frequency topographies of the interior VSH expansion of the discretized magnetic field ***B*** (see derivation and discussion in Section 2). Increase in *L* corresponds to an increase in spatial complexity.

One challenge in estimating the spatial frequencies present in the data is that the higher spatial frequencies decay faster as a function of distance [34]. Thus, the MEG signal measurement ***ϕ*** often appears spatially smooth at the sensor location, because lower spatial frequencies dominate. Resolving higher spatial frequencies is important for disambiguating the true multiple dipole source configurations, because different configurations may have the same lower order topography and only differ in their higher order spatial topography. New and improved methods of basic component extraction, especially at higher spatial frequencies, are therefore vital for accurately recording and reconstructing brain responses from MEG sensors.

In OPM-MEG systems, the sensors are placed much closer to the head, and therefore closer to the brain activity of interest. This gives OPMs a better opportunity to capture the higher-frequency components of the underlying neuronal activity, and therefore improve sensitivity and source localization [42], [34], [4]. However, increasing the number of spatial frequency components kept in the SSS basis can introduce instability and hinder the ability to separate the internal signals from the external magnetic fields, especially in cases with high sensor noise and MEG systems with fewer sensors [34]. In order to leverage the potentially improved spatial resolution provided by on-scalp MEG sensors, data preprocessing methods built on the basic component extraction of the measured magnetic field orders need to be optimized to account for instability, particularly with sensor noise and artifacts.

Some existing methods, such as the iterative SSS method, improve the stability of the SSS reconstruction by exploiting the hierarchical nature of the VSH expansion to iteratively process the data for each spatial component order [18]. Other variants of the SSS methods, such as the mSSS method and adaptive multipole models, adapt the geometry of the VSH expansion in SSS to on-scalp MEG systems, but these variants do not explicitly account for impacts from sensor noise [27], [38]. Spatiotemporal SSS (tSSS) incorporates a time-domain correlation analysis step that aims at recognizing and suppressing any signal components that are not satisfactorily suppressed by spatial SSS [35]. The better the spatial model is, the more robustly tSSS works in removing artifacts arising from, e.g., sources that are very close to the sensors. In this paper, we address improvements for spatial SSS and will not investigate tSSS further. Other statistical methods, such as independent component analysis (ICA) and signal space projection (SSP), utilize empty-room recordings or baseline periods to project out unwanted signals and noise, but once again, do not directly calculate or account for sensor noise itself, and require empty-room recordings for optimal performance [22], [37]. Obtaining accurate empty room measurements is paramount to avoid the risk of including brain activity in the noise projection, but is more difficult with OPM-MEG as the sensors move to dynamically fit the participant’s head, so they will not be in the same location during an empty room recording with no participant present.

As the reconstruction of brain signals using SSS methods requires the inverse of the spatial matrix, **S**, the task of reducing noise in the higher components becomes a linear algebra problem of matrix inversion to find solutions to a simultaneous linear equation. Here, we explore a novel application of the Wiener-Kolmogorov smoothing theory to matrix inversion called Foster’s inverse, after the author Manus Foster [13]. We evaluate the method’s ability to reduce noise when applied to the reconstruction of MEG data,with both cryogenic and on-scalp systems. Across simulations and MEG recordings, we compare this novel methods performance to other common MEG preprocessing methods: hihg-pass filtering, SSP, traditional SSS, and iterative SSS. Foster’s inverse utilizes the sensor noise covariance of the MEG system to weight and stabilize the SSS method, allowing for accurate reconstructions of the internal brain activity in the face of low frequency fields, large sensor noise, and artifacts.

## 2. Theory

### 2.1 Signal Space Separation Method and Variants

The Signal Space Separation (SSS) method decomposes the MEG signal vectors into two expansions of spherical harmonic functions, one that contains sources within the sensor helmet and one that contains sources outside the sensors, to separate brain activity from external magnetic fields [36],[34]. The origins of the electromagnetic signals measured by the sensor array can be deduced based on how the signals decay. Signals arising from sources within the brain should not decay at the origin, while signals that come from sources located outside the helmet should. We can further restrict the plausible subspace by only looking for magnetic signals that obey the quasistatic approximation of Maxwell’s equations. In this sense, the magnetic field **B** can be derived from the harmonic scalar potential *V*, and can be separated into two different sets of spherical harmonic basis functions, which are linear combinations of solutions to Laplace’s equation.

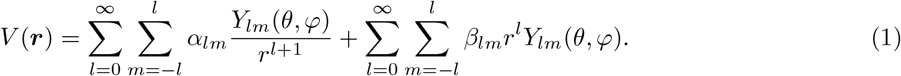

Where *Y*_*lm*_(*θ, φ*) is the normalized spherical harmonic function related to the associated Legendre polynomials and the radius *r* is the distance from the origin to the outside of the head. The first term of the sum corresponds to signals that originate inside the brain and decay at large *r*, whereas the second term represents signals that are stronger at large distances away from the origin and thus come from signals external to the brain. *α* terms denote internal signals (see Appendix 7.1 for further derivation), *β* terms denote external signals, and *V* is the total harmonic scalar potential such that [34]

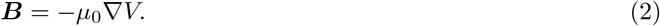

From the spherical harmonic discretization of *V*, the magnetic field ***B*** can also be calculated as an expansion of vector spherical harmonics (VSH)

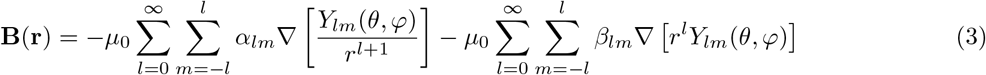

where multipole moments *α*_*lm*_ and *β*_*lm*_ weight each topographic VSH orders *l* and *m*. The full form of the multipole moments can be found in Appendix 7.1. After defining the radius *r* as the distance between the center of the brain and the closest MEG sensor, the VSH expansion of the magnetic field has two distinct terms, where the first represents brain signals that decay at large distances and the second term represents magnetic signals arising from outside of the brain [34]. This basis separation assumes the quasistatic approximation, such that Equation 2 is valid, and that the sensors are located in a current-free volume outside of the interior brain space [16], [22].

For computational applications, the measured MEG signal vector ***ϕ*** can be written as follows

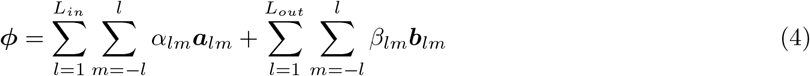

where ***a***_*lm*_ and ***b***_*lm*_ are the signal vector responses of the multichannel measurement device and ***ϕ*** = [*ϕ*_1_…*ϕ*_*N*_]^*T*^ is the measured signal vector across all *N* channels. The *l* = 0 components are excluded because they correspond to magnetic monopoles, which do not exist according to Maxwell’s equations. This form of ***ϕ*** can be expressed in matrix notation as:

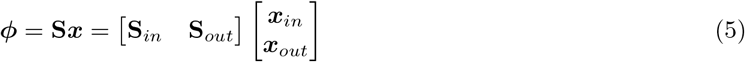

Where **S** is the magnetic subspace containing separate subspaces for internal and external sources, and the corresponding multipole moments are contained in vector ***x***. The moments *L* of the interior and exterior expansions are truncated at *L*_*in*_ = 8 and *L*_*out*_ = 3 [36], resulting in a (*L*_*in*_ + 1)^2^ + (*L*_*out*_ + 1)^2^ −2 = *D* dimensional space, where **S** is *N* MEG sensors by *D* in size (D = 95 in this case). The interior basis represented by **S**_*in*_ can be used to reconstruct the raw data in a way that excludes any noise that came from signals outside of the helmet, allowing for a clearer and more accurate representation of the signals that were produced by the brain during a specific period of time. The SSS method and Maxwell Filtering protocol are implemented in MNE-Python [24].

In principle, the signal vector ***ϕ*** is a linear superposition of in and out and noise vector ***n*** (*N* × 1 dimension), not explained by model of brain signal and outside world. The covariance of this noise, **N** (*N* × *N* dimension), can contain sensor noise and other nearby artifacts that result in noise in the higher spatial frequency range. We can represent the signal vector ***ϕ*** with noise as

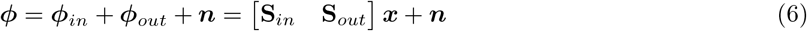

where an estimate for the multipole moments 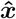 can be calculated by taking the direct psuedoinverse of the **S** matrix, **S**^†^, including the noise,

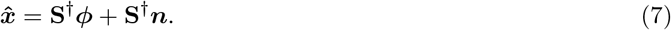

With the direct inverse, there is no way to avoid multiplying ***n***, which can explode the noise of the reconstructed data depending on the condition number and properties of the **S**. Specifically, sensor noise impacts the estimate of the multipole moment vector through the covariance 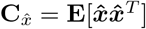 as [34]

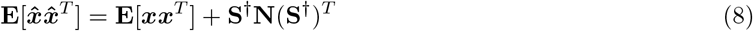

where **E** is the expectation value. Here, we see that the covariance of the sensor noise **N** directly impacts the estimation of the multipole moments, and in turn the reconstruction of the data using SSS. To avoid detrimental impacts from noise, other inverse methods such as iterative SSS have been proposed to improve the reconstruction of noisy data [18]. The iterative method modifies the estimate of the multipole moments by separating the inner subspace into distinct orders 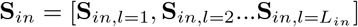 such that each multipole moment weight for each order can be estimated using only the corresponding partial **S**_*in*_ basis. For example, the first component is estimated as

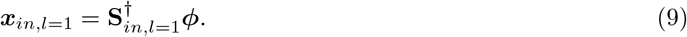

After iterating through each order, the resulting estimate for the weights ***x***_*it*,*in*_ can be used to reconstruct the internal measured signals using the same method as before: ***ϕ***_*in*_ = **S**_*in*_***x***_*it*,*in*_ [18]. Other adaptations of SSS include the spatiotemporal SSS method (tSSS) designed to reduce artifacts from head movement and nearby sources [35], and the extended SSS method (eSSS) designed to lower detrimental impacts from geometry and sensor calibration errors with a statistics based model built with SSP [17]. However, these methods do not directly account for the sensor noise itself mathematically, and may be unstable or cause the spreading of artifacts from one channel to another.

### 2.2 Presentation of Foster’s Inverse

Given some data vector ***y*** can be written ***y*** = **A*z*** + ***η***, where ***η*** is the noise vector and ***z*** is the solution or signal vector being estimated. We can solve for the Foster’s inverse estimate of the signal vector 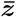 by performing the inverse operation [13]

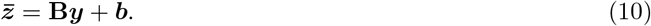

The task is to determine the matrix **B** and signal vector ***b*** such that we get the optimal estimate of the signal vector 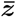. The solution depends on what is known about the noise and signal vector. If we assume that the noise and signal vector are independent, then the information needed is the two covariance matrices **Z** and **N** made up of the signal and noise vector components, respectively.

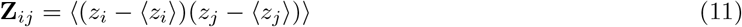

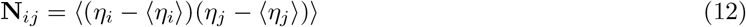

Using these matrices, an optimum inverse **B** of matrix **Z** is given by the Foster’s inverse

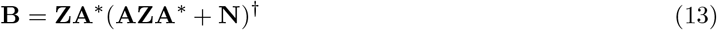

which uses the Penrose-Moore pseudoinverse. In fact, Foster’s inverse generalizes to the Penrose-Moore psuedoinverse in the absence of noise [13]. An optimal estimate of ***z*** solution vector is given by

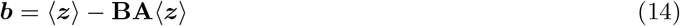

such that ***b*** is constant over the time course of the data collection. As such, Foster’s inverse can be applied to any scenario or model involving the relationship ***y*** = **A*z*** + ***η***. In fact, models with this form often appear in noise-normalized MNE methods for source localizations, with parallels and potential consequences discussed in Appendix 7.2. But for the purposes of this paper we will focus on adapting Foster’s inverse with the SSS method as motivation.

### 2.3 Adaptation of Foster’s Inverse with the SSS Method

In the context of MEG and SSS, Foster’s inverse offers an alternative to the multipole moment estimate in Equation 2.1, allowing for a more robust estimate utilizing the sensor noise covariance. First, let ***y*** be the measured signal vector ***ϕ***, and matrix **A** be the VSH basis obtained from SSS, matrix **S**. In principle, the matrix **S** can be obtained from other SSS variants [27],[38], as long as the corresponding multipole moments are calculated to reflect the change in basis representation. Specifically, we implement Foster’s inverse with the traditional SSS basis and the mSSS basis, denoted matrix **S**_m_, as this method has been optimized for adapting SSS to on-scalp MEG systems [27]. Next, the solution vector ***z*** is the multipole moments ***x***, which will be estimated given the SSS basis and the MEG data.

Then, the last adaptation in the Foster’s inverse problem is defining ***η*** as the MEG sensor noise. Other sources of magnetic interference are classified as signal, whether these sources pertain to brain activity or not, and are represented in the SSS matrix by design. We want to write the signal vector ***ϕ*** using the SSS basis, multipole moment coefficients, and noise as described in Section 2

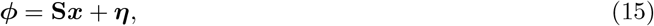

where we want to optimize the inverse equation

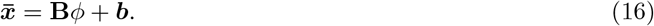

Matrix ***B*** is Foster’s inverse of the SSS matrix ***S***, [13]

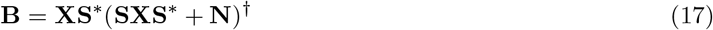

and

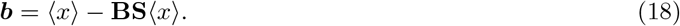

Then, the improved estimate 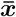 can be used to reconstruct the interior signal in the same fashion as traditional and iterative SSS.

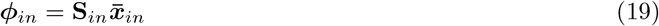

The matrix **X** is determined using the covariance of the multipole moments found using SSS as

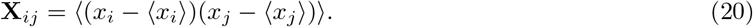

Alternatively, if the location of the current dipole source is exactly known, perhaps in a forward model calculation or otherwise, the multipole moment coefficients *α*_*lm*_ can be calculated directly using Equations 25, 26, and 27 as presented in Appendix 7.1.

The noise covariance matrix **N** can be estimated, calculated, or measured. It must correspond to sensor noise, so any magnetic signal features in matrix **S** should not be included in the noise matrix. In the case where the sensor noise covariance is unknown, Foster’s inverse can be applied in a case of minimum information where the noise-to-signal ratio is used as a proxy for the full covariance profile, the full derivation of which can be found in Appendix 7.3. Here, we focus on two methods for calculating **N**: the empirical method and the over-sampled temporal projection (OTP) method, described in detail in the next section (Section 3).

## 3 Sensor Noise Covariance Profile Estimation

By definition, covariance is the measurement of how two random variables change together. In the context of MEG measurements, the sensor noise covariance describes the spatial covariance of noise between sensors and other artifacts that are not captured in the external expansion of the VSH basis, which accounts for sources of external interference. There are a variety of ways to quantify or estimate the level of noise in the sensor array, which impacts the noise covariance profile used in data processing steps such as beamformers, MNE dipole localizations, and estimates in quantifying the impacts of noise in SSS[24], [22], [36]. The method for determining the sensor noise covariance of an MEG system also determines the performance of preprocessing with Foster’s inverse and SSS. The goal with our SSS-based implementation is to obtain a measurement of the noise covariance without including any magnetic signals that fit a distinct spatial model, such as VSH, which are already characterized by the external components of the SSS matrix. The noise covariance profile for the sensors includes any noise that is unique to each sensor itself, perhaps due to differences with manufacturing, as well as any interactions between sensors from spatially correlated noise which may be from fluctuations in sensors themselves or some magnetic impurity near the sensors that effects multiple sensors at once.

The simplest noise covariance profile is a diagonal matrix which represents a case where there is no spatially correlated noise between the sensors, where purely sensor related noise is assumed to be independent across sensors [10]. Here, the external SSS basis is needed to capture other sources of interference and separate them from the internal brain signals. In practice, however, off-diagonal elements occur, which has been widely characterized for SQUID-MEG sensors corresponding to spatial correlation in noise resulting from hardware specifications of the sensors, and occurs in OPM-MEG sensors due to quantum, optical, and magnetic effects [43], [40]. Noise covariance profiles that are not assumed to be purely diagonal (not containing spatially correlated noise) must be estimated. Practically, some methods for measuring sensor noise involve taking an empty-room recording in the magnetically shielded room (MSR) prior to testing, and then calculating the noise covariance from that recording [24]. A similar result can be obtained by using resting-state segments of a recording, or from pre-stimulus baselines, which should include minimal brain activity.

Here we test two methods for calculating the noise covariance matrix implemented with Foster’s inverse: 1) the empirical covariance estimation [10], and 2) a novel application of the Oversampled Temporal Projection (OTP) method, and we describe the considerations and applications of both methods in the following sections [23].

### 3.1 Empirical Covariance with MNE-Python

MNE-Python is a powerful package for analyzing MEG data, which implements several functions for estimating the noise covariance from raw and epoched MEG data [24]. The majority of the different methods for calculating noise covariance are characterized by Engemann and Gramfort, and assume that the amplitudes of the sources and measured data are Gaussian due to linear mixing, therefore the additive noise is also assumed to be Gaussian such that the total MEG signal can be characterized with the combination of a mean vector and the corresponding covariance matrix [10]. Here, we focus on one of these presented methods called the empirical covariance, which is the default method implemented in MNE-Python.

The empirical covariance matrix **N**_E_ of data matrix **Y**, dimensions *N* × *M* can be calculated as 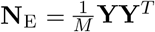. If the number of data points *M* is very large, the empirical covariance model becomes a better estimate of the true covariance, so the empirical noise covariance should be calculated on longer periods of baseline recordings or empty-room recordings. However, in the absence of empty room recordings, the number of baseline periods for noise estimation during data collection may be too small to obtain a good estimate, or the estimate may include brain signals in the noise.

For the implementation of Foster’s inverse using the empirical method of noise covariance, in cases where an empty-room recording is available, the covariance is estimated from the empty-room. In the absence of an empty-room recording, the initial baseline period before stimulation is used to estimate the noise covariance. In datasets with both empty-room recordings and not, we use only the internal SSS basis (containing the first 80 expansion components) with the empirical covariance in Foster’s inverse as the external basis and estimate for the noise covariance may contain overlapping information [36]. Additionally, the noise covariance in the sensors may be impacted by the fields in the external basis, especially in the case of OPM-MEG systems where the sensors are internally modulated and compensated based on the fields in the room [1], [4].

### 3.2 Oversampled Temporal Projection to Calculate Noise Covariance

Alternative methods for reducing the effects of sensor noise within MNE-Python, such as the Oversampled Temporal Projection (OTP), only require that the signals of interest are spatially oversampled by the number of sensor channels and that the signals of interest are statistically independent from the noise [23]. OTP is able to capture random sensor noise fluctuations and remove them from the data. In order to do this, we first define the total data matrix **D**, which is composed of *N* sensor channels with data over *n*_*s*_ temporal samples. The data matrix can be expanded as:

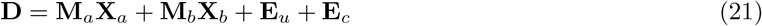

where **M**_*a*_ defines the subspace for signals of interest and **M**_*b*_ defines the subspace for external interference signals, and **X**_*a*,*b*_ define the respective coordinates. Matrix **E**_*u*_ contains uncorrelated artifacts and individual sensor noise, whereas **E**_*c*_ contains correlated artifacts that are set to zero and not included in the OTP model [23]. We can then combine this equation into a compact matrix equation

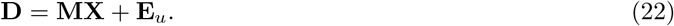

Where the matrices **M** and **X** contain both the concatenated subspaces *a* and *b*. Finally, the noiseless signal **D**_0_ can be calculated as

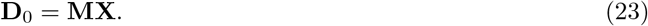

The difference in the raw data and the OTP processed data is the effect of the sensor noise. The covariance of the sensor noise can then be calculated as **N**_OTP_ = covar(**D** −**D**_0_), and can then be used in Foster’s inverse. Unlike the empirical method, the OTP method for estimating noise isolates random sensor noise, and is implemented in conjunction with the full SSS basis in Foster’s inverse.

## 4 Methodology

### 4.1 Implementation of Foster’s Inverse

We implemented the novel methodology of Foster’s inverse with SSS in Python [25], and integrated it with MNE-Python functionality for calculating the signal space separation (SSS) basis and the noise covariance matrix using both the empirical and OTP methods [24], [23]. We also implemented Foster’s inverse in MATLAB for the purpose of preliminary simulation investigations, before moving to Phantom and subject MEG data processing in Python. As described in Section 3, the choice of method for computing the sensor noise covariance matrix influences the choice for the SSS basis used in Foster’s inverse. The empirical noise covariance, **N**_*E*_, is used in conjunction with the internal SSS basis when it is estimated from an empty-room recording and from a baseline period. The OTP method is implemented with the full SSS basis.

We also test two different variations of Foster’s inverse in two different case studies to show that the mathematical method is robust to different methods of obtaining the spatial basis matrix and the sensor noise covariance matrix, which may change for different MEG systems and scenarios. First, the novel OTP method for calculating the sensor noise covariance is compared to the empirical method in Section 5.3. As the OTP method is designed to isolate random sensor noise, the resulting covariance matrix **N**_OTP_ is used with the full SSS basis because the noise covariance does not contain any information about the external magnetic fields. Next, we implement Foster’s inverse with the multi-SSS (mSSS) method in Section 4.5.3, an adaptation of SSS specifically modified for on-scalp MEG systems like OPM-MEG [27].

### 4.2 Comparison Metrics

We calculate a number of comparison metrics to investigate the performance of high-pass filtering, SSP, SSS, iterative SSS, and Foster’s inverse with SSS on a variety of different datasets. As we are primarily interested in noise and sensor noise, the first metric is the signal-to-noise ratio (SNR) calculated from a baseline period of no brain activity (noise) compared to the peak brain activity (signal). The SNR for each dataset is calculated as:

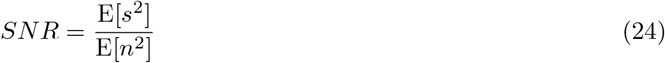

where the expected value E is the mean square, or mean of the standard deviation squared, equivalent to the mean of the variances, of the signal *s* or noise *n*. This method is used for both simulations and for single-subject MEG recordings. This metric is implemented across all the data types, and each data and MEG system type has its own unique comparison metrics which will be described in the following sections.

### 4.3 Current Dipole Simulations

Here, we implement a current dipole simulation in MATLAB with a single current dipole located at 5cm along the x-axis using the Sarvas formula [33]. The signal is generated for 0.5 seconds with a timestep of 0.001 seconds, and the moment of the dipole varies sinusoidally in time. Before the dipole is activated, the simulation includes a baseline period to represent the time when a brain is less active before experiencing the onset of a stimulus, and Gaussian noise is added at a level of 20% of the maximum simulated signal amplitude. The random seed for generating this noise is held constant, therefore there is no need for multiple trials per simulation. Finally, the magnetic flux through each of the sensors was calculated using standard gradiometer and magnetometer dimensions for the 306-channel MEGIN/Elekta Neuromag SQUID sensors given in the MNE-Python documentation [24].

Because the simulations were performed in MATLAB, Foster’s inverse was also implemented in MATLAB here. For the implementation of Foster’s inverse, optimizing the multipole moments 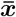 is more straightforward in simulations because the initial multipole moments ***x*** can be calculated directly using the location and moment of the simulated dipole up to the internal expansion order *L*_*in*_ as described in Appendix 7.1. We then calculate the diagonal noise covariance matrix **N** from the generated Gaussian noise. Foster’s inverse is compared to SSS and iterative SSS using SNR and power-spectral density (PSD).

This simulation serves as a proof-of-concept for the mathematical method, where we can perfectly calculate the SSS basis, the corresponding multipole moments exactly from the simulated dipole, and the noise covariance from the exactly known simulated sensor noise. In the following applications of Foster’s Inverse with SSS to real MEG data, the multipole moments and the sensor noise must instead be calculated and estimated.

### 4.4 Phantom Head Recordings

The dry Phantom head manufactured by MEGIN/Elekta Neuromag comes with the SQUID sensor system as a diagnostic tool. The head is made of a plastic hollow sphere with radius 87.5 mm containing 32 artificial current dipoles that can be activated independently in series with four HPI coils for head position tracking. The current dipoles themselves are made using equilateral triangles of current lines to produce a magnetic field distribution equivalent to a current dipole oriented tangentially to the surface of a spherical conductor [22], [9]. Since the orientations and locations of the 32 dipoles are exactly known, the data measured by the SQUID-MEG system when each dipole is activated can be treated as an evoked set and used to check the localization of the dipole.

Here, we focus on four different Phantom datasets taken with the 306-channel SQUID sensor MEGIN/Elekta Neuromag MEG system. These datasets were taken at the Institute for Learning and Brain Sciences (I-LABS), University of Washington, between 2023 and 2024, with varying dipole field strengths, reflecting the input current or voltage levels, and internal active shielding (IAS) settings. The file names listed in Table 2 contain the exact date as year-month-day, followed by the dipole field strength relating to the current passed through the wires, and the IAS system either on or off. The datasets are numbered 1) through 4) in the first column for ease of reference. Datasets 1) and 2) were recorded on the same day with two different dipole field strengths, 1000 nam and 200 nam, respectively. Similarly, datasets 3) and 4) are from the same day, both using 1000 nam levels in the current dipoles, with the IAS system off, then on. 1000 nam is unrealistically strong, whereas 200 nam is a better approximation of human brain activity.

Each dataset was processed with traditional SSS and Foster’s inverse using the empirical method of calculating the noise covariance **N**_E_ implemented in MNE-Python. The Foster’s inverse calculation is implemented in Python for the Phantom analysis, whereas the simulated results discussed in the previous sections were implemented in MATLAB. As the iterative SSS method is only implemented in MATLAB, we focus on only the SSS method compared to Foster’s inverse here. Each of the 32 dipoles is activated multiple times, and the resulting data is averaged into epochs, which are then used to localize the dipole location in MNE-Python [24]. To quantify the success of the dipole localization calculation, we calculated the mean and maximum error in position and orientation across all 32 dipoles, as well as an overall goodness of fit (GOF) percentage where a higher GOF indicates the dipole localization was the most accurate across each of the 32 Phantom dipoles, accounting for both position and orientation.

### 4.5 OPM-MEG Recordings

We verified the ability of Foster’s inverse with SSS, implemented in Python, to accurately reconstruct the peak brain activity while mitigating the impact of sensor noise using single-subject datasets from two different OPM-MEG sensor systems. All data were collected in accordance with the given university’s institutional review board (IRB) human subjects testing guidelines, and informed consent was obtained from all participants for being included in the study.

#### 4.5.1 Single-Subject Kernel Flux OPM-MEG Recording

The first single-subject OPM-MEG dataset was collected at I-LABS at the University of Washington (UW) with the 432-channel Kernel Flux OPM system with coil sensing directions oriented perpendicular to the surface of the subject’s head [26]. The subject listened to 180 repeats of a single frequency tone. Evoked data windows of approximately one second long averaged over 180 trials of the audio tone response are used for analysis with baseline correction applied [24]. The raw data are first high-pass filtered at 0.1 Hz and low-pass filtered at 40 Hz. Then, the raw data are preprocessed with SSS, Iterative SSS (preprocessed using MATLAB, then visualized using MNE-Python), and Foster’s inverse using two different methods of calculating the noise covariance: the empirical method (**N**_E_) with the internal SSS basis and the novel OTP (**N**_OTP_) method application with the full SSS basis, both done using MNE-Python. The empirical covariance is calculated from an empty-room recording, and the OTP method is applied to the raw data. We compare the performance of each preprocessing method by calculating the SNR and visualizing the evoked butterfly plots and topographies at the peak time of activation.

#### 4.5.2 Multi-Subject FieldLine OPM-MEG Recordings

Second, we collected six single-subject datasets using the 144-channel single-axis SERF FieldLine OPM system at Stanford University. Each subject listened to 200 repeated trials of a 400 Hz audio tone lasting for 1 second, with 1.2 seconds between each tone onset, generated and delivered using PsychoPy [31]. We first focus on a single-subject case study exhibiting one highly noisy sensor, then compare evoked responses across all six subjects with different preprocessing methods: high-pass filtering, SSP, traditional SSS, and Foster’s inverse with SSS using the empirical method for noise covariance calculation (**N**_E_). Because empty-room recordings with the OPM sensors cannot be measured with the sensors in the same position with and without a participant, the noise covariance was estimated using the baseline period before stimulation, and is used in conjunction with the internal SSS basis in Foster’s inverse. Coregistration between the sensor locations and MRI space was achieved using the RevoPoint 3D structured light scanner and the YORC pipeline developed at the University of York [2], which was used to calculate the transformation matrix between the OPM sensor system and the subject MRI head space as defined by and implemented in MNE-Python [24]. For three subjects with individual structural MRIs, we proceed with forward and inverse modeling for each preprocessing method after the raw data is filtered between 0.5-80 Hz and notch filtered at 60 Hz. First, the T1 MRI scans are segmented and processed using FreeSurfer and the MNE watershed algorithm to extract the desired cortical surfaces [11], [24]. The forward solution is made using MNE-Python with a three-shell conductivity model and source spacing set to ico4 [24]. The inverse operator is calculated using dynamic statistical parameter mapping (dSPM) for each preprocessing method with unsigned free orientation dipoles and no depth weighting to preserve the orientation information of the dipole sources [14] [24], [7]. dSPM is a noise-normalized linear estimate, the underlying mathematical method of which compliments the linear algebra techniques employed in Foster’s inverse as explored in Appendix 7.2 [7]. The results of dSPM are normalized dipole strengths at each location reported in Amplitude (AU), where the amplitude of the strength is defined against the noise floor null hypothesis comparable to an F-statistic, where larger values indicate better localization results [7]. For each subject and preprocessing method, we show the source localization at the time of peak activation and calculate the explained variance (EV%) of the peak signal compared to the dataset. Finally, dipole fitting is performed using MNE-Python, where one source is fit for each point in time during the evoked average with no minimum depth requirement and freely oriented dipoles, and we calculate the minimum and maximum dipole GOF (%).

#### 4.5.3 Foster’s Inverse with mSSS

Finally, we investigate the performance of Foster’s inverse with mSSS compared to with SSS, and with traditional SSS itself. The mSSS method is an adaptation of SSS for use with on-scalp MEG systems, thus we test Foster’s inverse with SSS on the FieldLine OPM-MEG system. We maintain a threshold of 5% significance, as described in the mSSS method implementation in MATLAB [27], and then visualize the preprocessed data in MNE-Python. Using the individual subject’s MRI, the mSSS basis is separately calculated with individually optimized expansion origins using a three-shell BEM model [24], [27]. First, we show averaged evoked responses with a high-pass filter at 0.5 Hz and a low-pass at 50 Hz. Then, the same steps were followed as described above in Section 4.5.2 to generate forward and inverse solutions. Next, for each subject we perform the same source localization steps as before to calculate the explained variance and goodness of fit percentages for the underlying current dipole sources, including visualization of the peak source localization. These procedures demonstrate how Foster’s inverse can be dynamically applied to a variety of different MEG sensor systems with unique considerations for calculating the spatial basis representing the signals.

## 5 Results

### 5.1 Single Current Dipole Simulation Results

For a proof of concept verifying the correctness of the method and its power when the covariance information is perfectly known, Foster’s inverse implementation of SSS is compared to the raw simulated signal, the traditional SSS reconstruction, and the iterative SSS reconstruction first by calculation of the SNR. The SNR calculation results using the baseline period before 50 ms as the noise period and the signal period after 50 ms are shown in Table 1.

**Table 1:** Signal to Noise Ratio (SNR)

| SSS Variant | SNR |
| --- | --- |
| Simulated Data | 6.08e+3 |
| SSS | 1.15e+4 |
| Iterative SSS | 1.10e+4 |
| Foster's SSS | 5.48e+5 |

Foster’s inverse provides over a tenfold increase in SNR compared to the iterative and traditional SSS reconstructions of the simulated data as evidenced in Table 1. These results are also seen visually when the simulated current dipole is plotted with the SSS-processed and Foster’s-processed data along the same axis.

As seen in Figure 2, the Foster’s inverse reconstruction of the simulated MEG data has vastly reduced the amplitude of noise compared to the unprocessed and SSS processed data. This result can be further seen in the PSD plots of the three datasets in Figure 3

**Figure 2.**
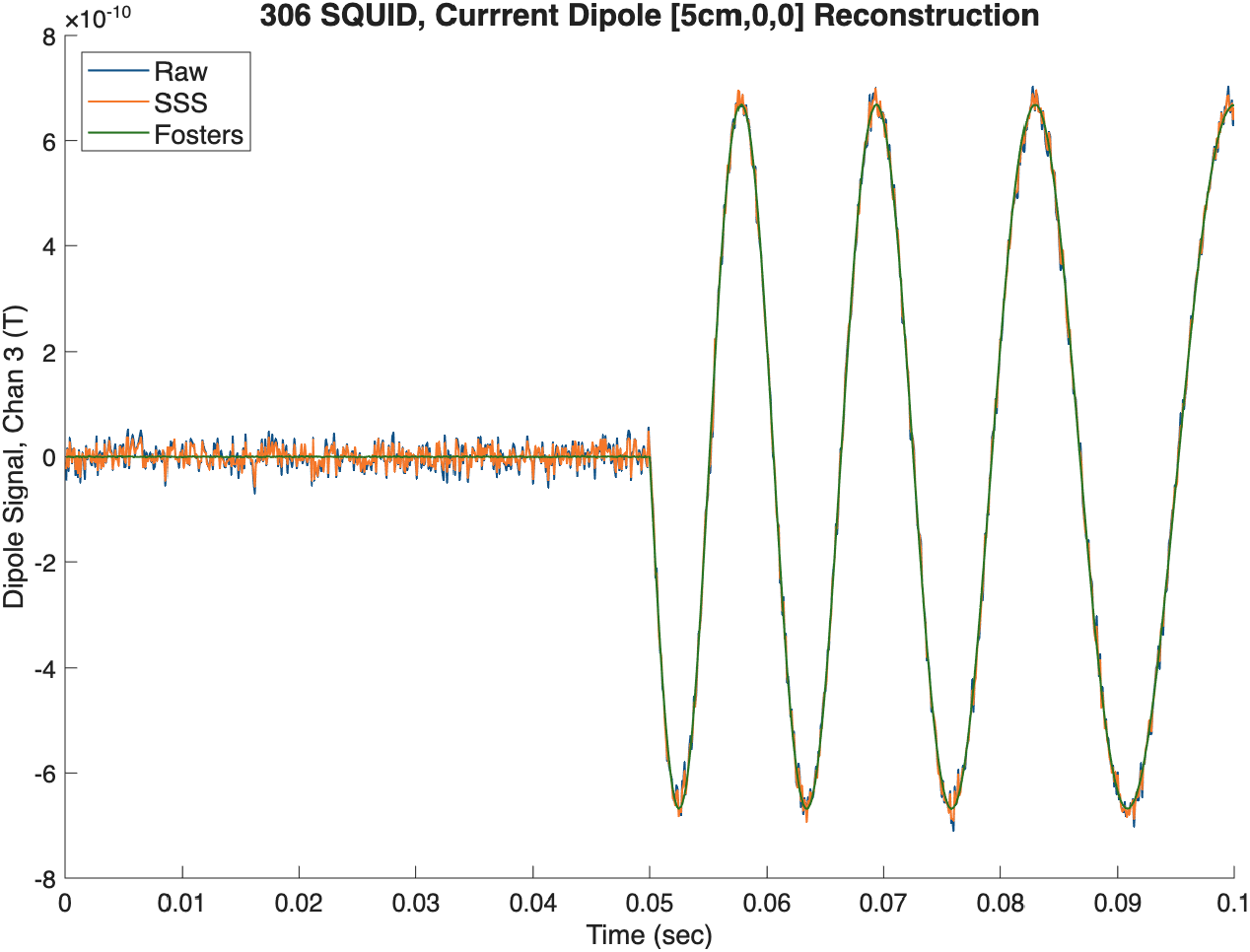
Comparison of raw noisy current dipole data and data after Foster’s inverse SSS reconstruction generated using the Sarvas current dipole formula with Gaussian noise added at 20% of the maximum signal.

**Figure 3.**
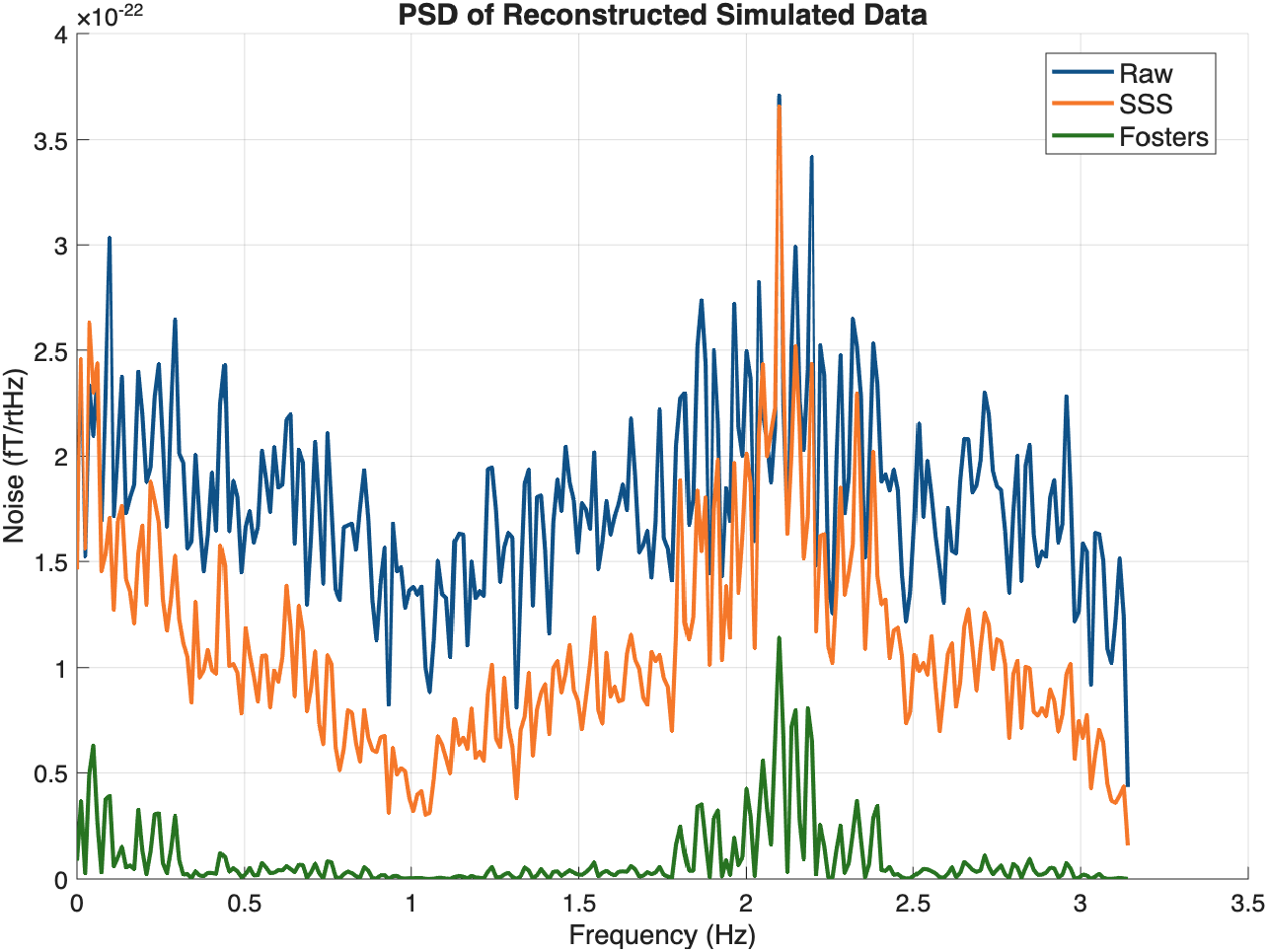
PSD plots of the simulated current dipole signal, SSS reconstructed signal, and Foster’s inverse SSS reconstructed signal. Plots are generated and calculated in MATLAB.

The dramatic decrease in signal reconstruction noise seen in Figures 2 and 3 is a promising result, suggesting the potential for the Foster’s inverse method to provide noise reduction on higher order terms when applied to both Phantom and human MEG data recordings. As mentioned, when the location of the dipole is known, the multipole moments can be calculated directly as in this simulation, and the simulated sensor noise is precisely known, leading to marked improvements in SNR. In reality, however, the location of the current dipole source in the brain is unknown, but can be estimated using the SSS reconstruction as described in Section 2, and the noise covariance must be calculated from the data, as we will implement and explore in the following sections.

### 5.2 Phantom Head Results

The results across all four Phantom datasets are shown in Table 2 and visualized in Figure 4, including the dipole localization error, the orientation error, and GOF metrics. Note that channels marked bad were not removed from the data before preprocessing.

**Table 2:** Mean and Maximum Dipole Localization Errors and GOF Metrics for Each Phantom Dataset.

| **Dataset** | Method | Position E (mm) | Orientation E (°) | GOF (%) |
| --- | --- | --- | --- | --- |
|  |  | mean/max | mean/max | mean/max |
| 1) 24-07-08,<br>1000nam | Raw | 2.8/12.2 | 3.3/16 | 83.0/99.8 |
|  | SSS | 10.5/58.2 | 9.4/48 | 86.7/99.8 |
| | Foster's $\mathbf{N}_E$ | 2.1/5.8 | 2.7/9.2 | 98.7/99.6 |
| 2) 24-07-08,<br>200nam | Raw | 14/103.4 | 14.8/81.1 | 63.4/97.9 |
|  | SSS | 21.6/98.0 | 24.2/84.7 | 73.0/99.1 |
| | Foster's $\mathbf{N}_E$ | 7.2/19.9 | 8.2/24.8 | 83.6/98.7 |
| 3) 23-12-07,<br>1000nam, IAS off | Raw | 2.1/4.2 | 2.4/6.3 | 98.3/99.9 |
|  | SSS | 3.5/9.0 | 3.4/13.6 | 98.3/99.9 |
| | Foster's $\mathbf{N}_E$ | 3.8/8.5 | 3.9/12.8 | 98.8/99.7 |
| 4) 23-12-07,<br>1000nam, IAS on | Raw | 2.9/7.9 | 3.4/13.5 | 98.4/99.8 |
|  | SSS | 3.1/7.1 | 3.3/9.7 | 99.2/99.9 |
| | Foster's $\mathbf{N}_E$ | 2.7/6.0 | 2.9/8.8 | 99.0/99.7 |

**Table 3:** Average Mean Dipole Localization Errors and GOF Metrics Across Each Phantom Dataset.

| **Method** | Position E (mm) | Orientation E (°) | GOF (%) |
| --- | --- | --- | --- |
| Raw | 5.45 | 5.98 | 85.8 |
| SSS | 9.68 | 10.1 | 89.3 |
| Foster's $\mathbf{N}_E$ | 3.98 | 4.43 | 95.7 |

**Figure 4.**
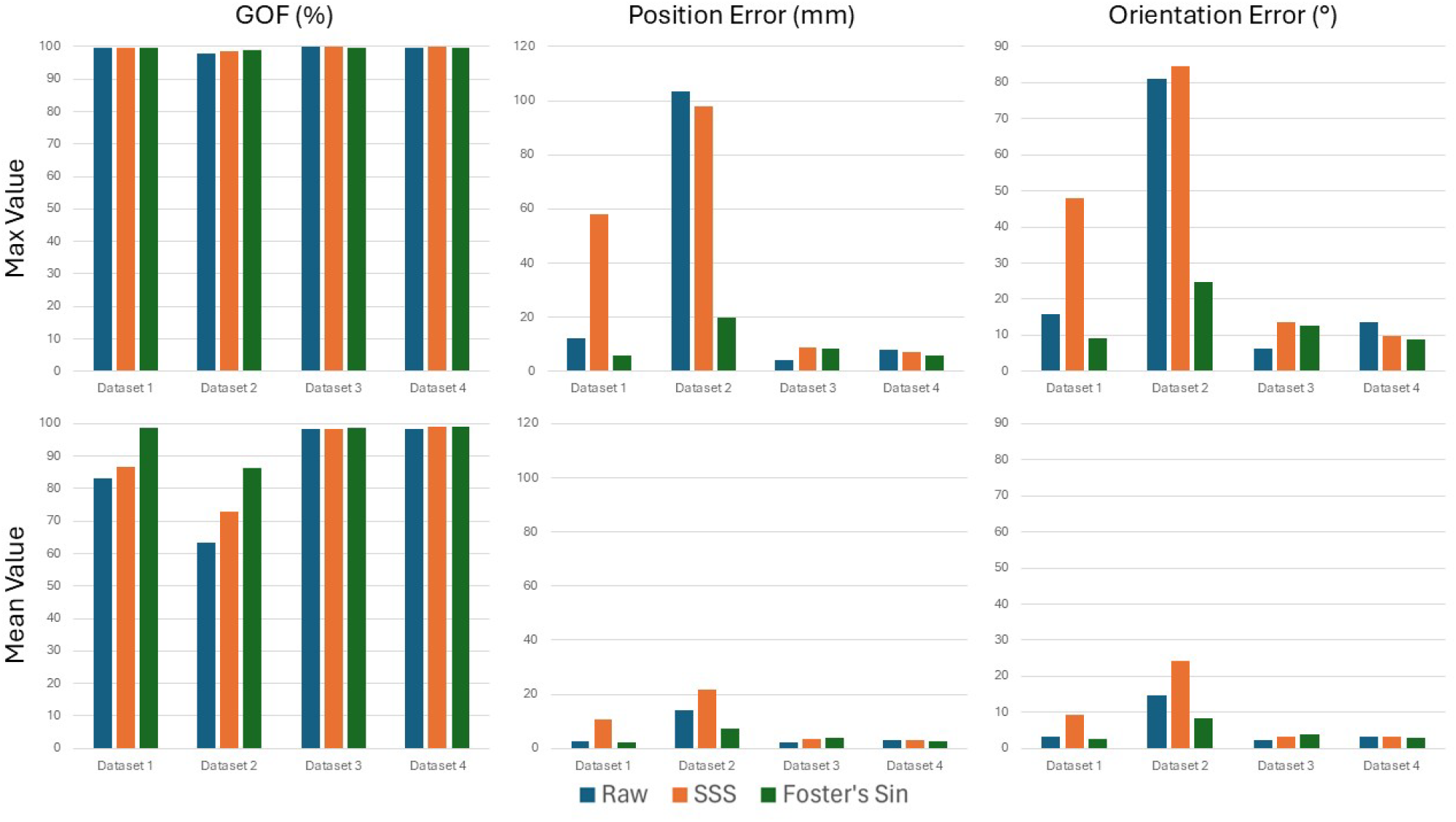
Mean and Max dipole localization and GOF metrics for each Phantom dataset, a visualization of the data in Table 2

Evoked plots of datasets 1) and 2) are visualized to showcase cases with some data abnormalities, as seen by the artifact specifically in the magnetometer channels in row a) of Figure 5 and 6. With no other artifact removal procedures, the SSS processed data does improve the GOF from an average of 83% to 86.7% (see Table 2). Crucially however, the method suffers from a spreading of the noise from the original artifact throughout other channels. As seen in row c) of Figure 5, Foster’s inverse with SSS is able to recover the expected dipole plots with a GOF of 98.7%. Similarly, in row c) of Figure 6, Foster’s inverse is able to recover the expected dipole oscillations despite the large magnetometer artifacts while the SSS reconstruction is not able to recover this, with the artifact also impacting the gradiometer channel reconstructions as well. Overall, as seen in Figure 4, Foster’s inverse either matches or improves the GOF of the dipole localization results, specifically by increasing the mean GOF and especially in datasets with larger noise and artifacts like datasets 1) and 2). Table 3 show that, overall and on average, Foster’s inverse with SSS reduced the dipole localization error by 2 mm and improved the goodness of fit percentage by 10%, showing that even in datasets with large artifacts, processing data with Foster’s inverse results in precise and accurate dipole localizations. In datasets without significantly apparent sensor noise and artifacts, such as datasets 3) and 4), Foster’s inverse with SSS and SSS perform comparably equally, with SSS decreasing the position and orientation error the most in dataset 3), but Foster’s inverse increasing the mean GOF across all dipoles.

**Figure 5.**
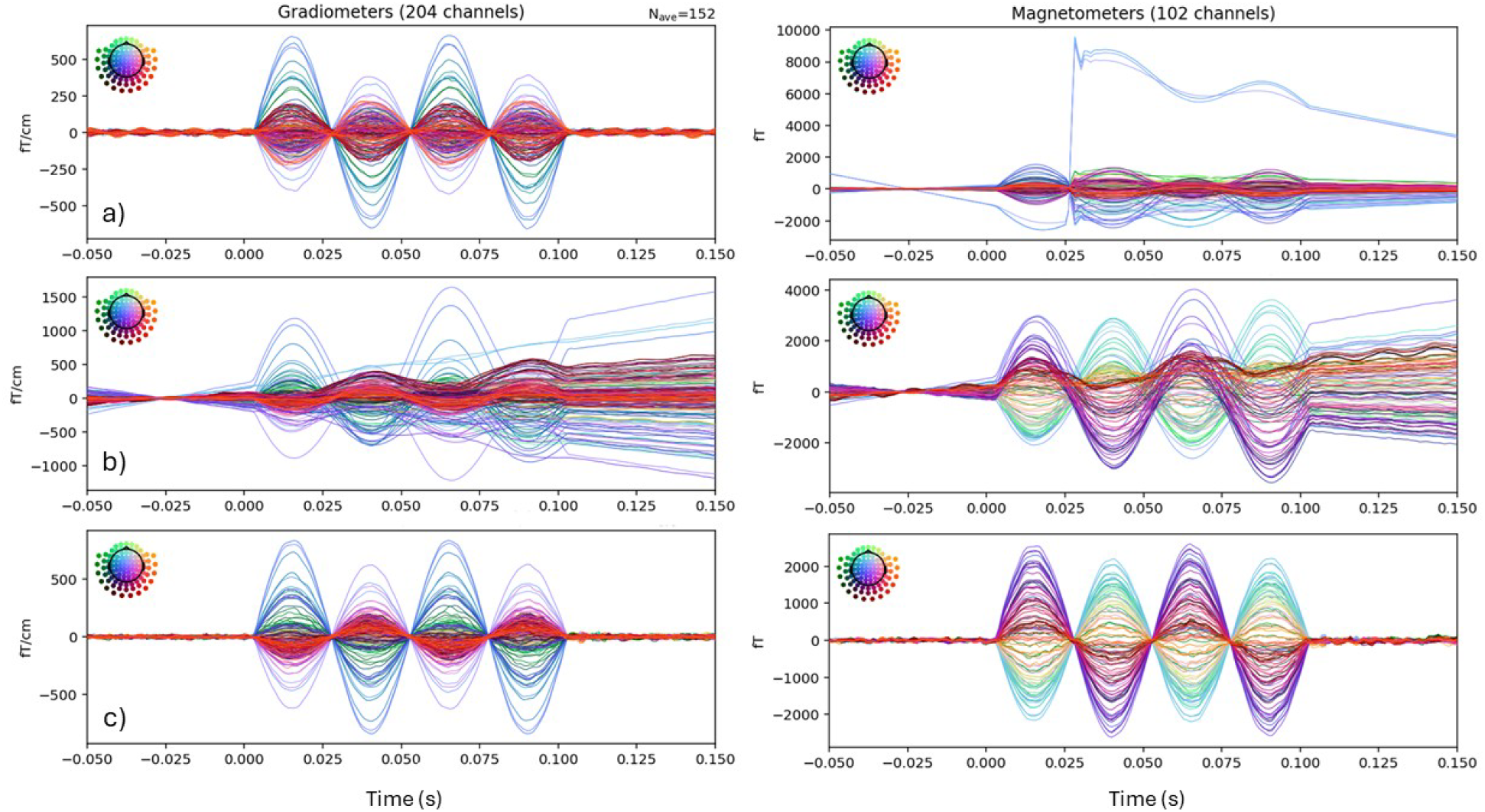
Butterfly evoked plots for Phantom dataset 1) generated using MNE-Python where row a) shows the unprocessed evoked data, b) shows the SSS processed data, and c) shows the Foster’s inverse with SSS processed data.

**Figure 6.**
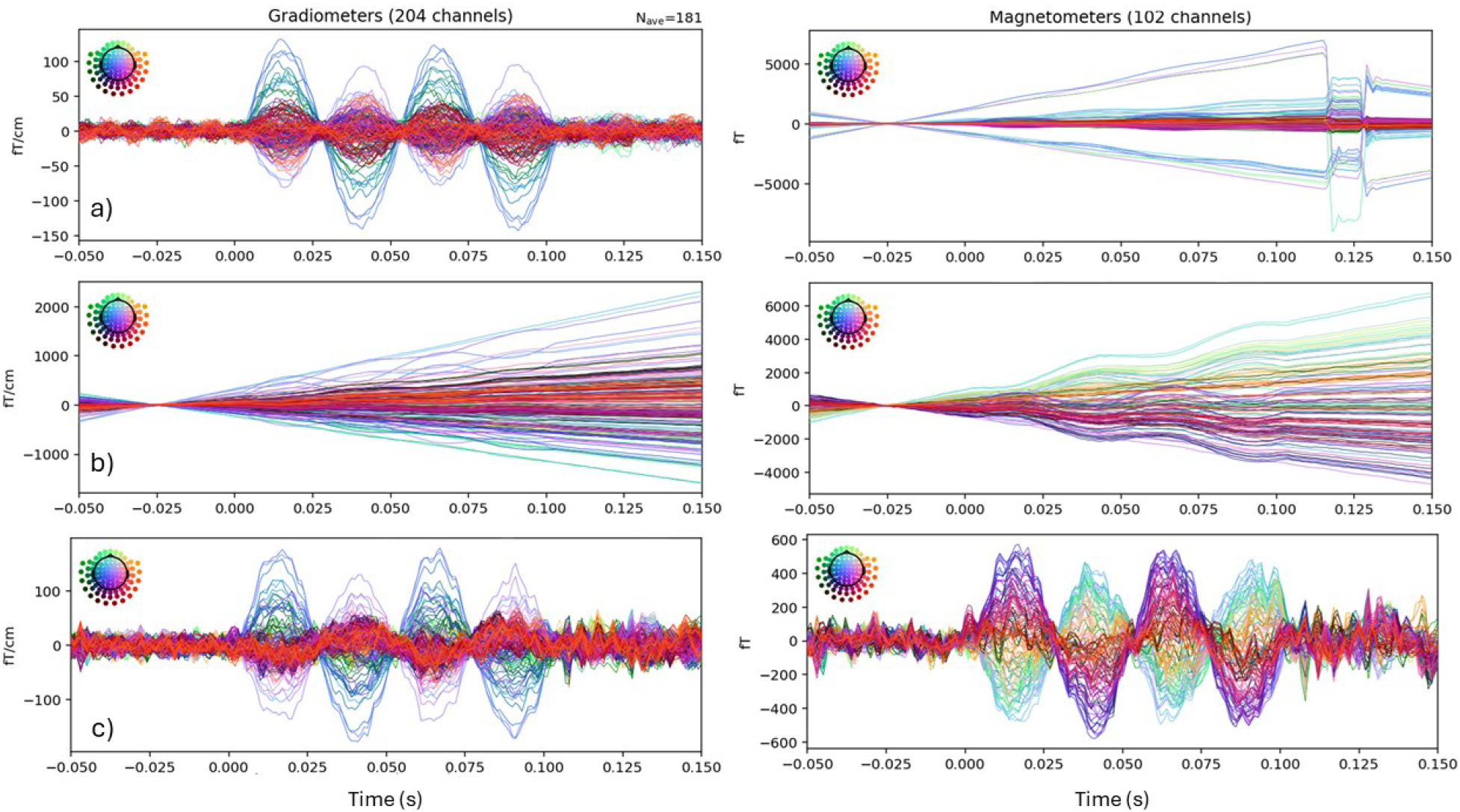
Butterfly evoked plots for Phantom dataset 2) generated using MNE-Python where row a) shows the unprocessed evoked data, b) shows the SSS processed data, and c) shows the Foster’s processed data.

General practice for some preprocessing methods, like SSS, recommends the removal of bad channels before preprocessing to avoid the spreading of artifacts like we have seen in Figures 5 and 6. In this example in datasets 1) and 2), these artifacts would be easy to uncover and the bad channels can be manually or potentially algorithmically removed, but in cases where the bad channels go undetected, Foster’s inverse with SSS mitigates the spread of the artifacts and allows for robust reconstructions of the brain signal.

### 5.3 Kernel Flux OPM Data

Here, we use the single-subject audio dataset collected using the 432-channel Kernel Flux OPM-MEG system. After preliminary analysis, 349 well-functioning channels were kept for analysis. The first evoked is preprocessed with a high-pass filter at 0.1 Hz and a low-pass filter at 40 Hz, the second is processed using SSS through the Maxwell Filtering function in MNE-Python, the third is processed using the iterative SSS method in Matlab with 10 iterations then transferred to Python for visualization, the fourth is Foster’s inverse with the internal SSS basis and **N**_E_, and the fifth is Foster’s inverse with the full SSS basis and **N**_OTP_ calculated using the MNE-Python implementation [36]. Butterfly plots of the evoked waveforms and topographic plots at the peak activity time are visualized for comparison, and the SNR is calculated in the same manner as described previously.

Along with SSS and iterative SSS, Foster’s inverse with **N**_E_ is able to reduce the noise levels in the data to clearly isolate the peak brain response. Foster’s inverse increases the SNR as seen in Table 4, corresponding to a visual decreased in the amplitude of noise during the periods before an after the peak response from 0.1 s to 0.2 s in as seen in Figure 7. Furthermore, Foster’s inverse with **N**_E_ and **N**_OTP_ recover the expected peak auditory evoked topographies as SSS and iterative SSS 8. However, Foster’s inverse using the OTP method for estimating the noise covariance under performs the empirical method in SNR. This may be because the sensor noise is not purely random, but contains other sources of noise impacting OPM-MEG sensors [32], [4] which are better captured by the empirical covariance method.

**Table 4:** Signal to Noise Ratio (SNR) of Kernel Flux OPM Evoked Dataset.

| SSS Variant | SNR |
| --- | --- |
| Filtered | 11.8 |
| SSS | 23.1 |
| Iterative SSS | 29.4 |
| Foster's $\mathbf{N}_E$ | 31.4 |
| Foster's $\mathbf{N}_{OTP}$ | 21.5 |

**Figure 7.**
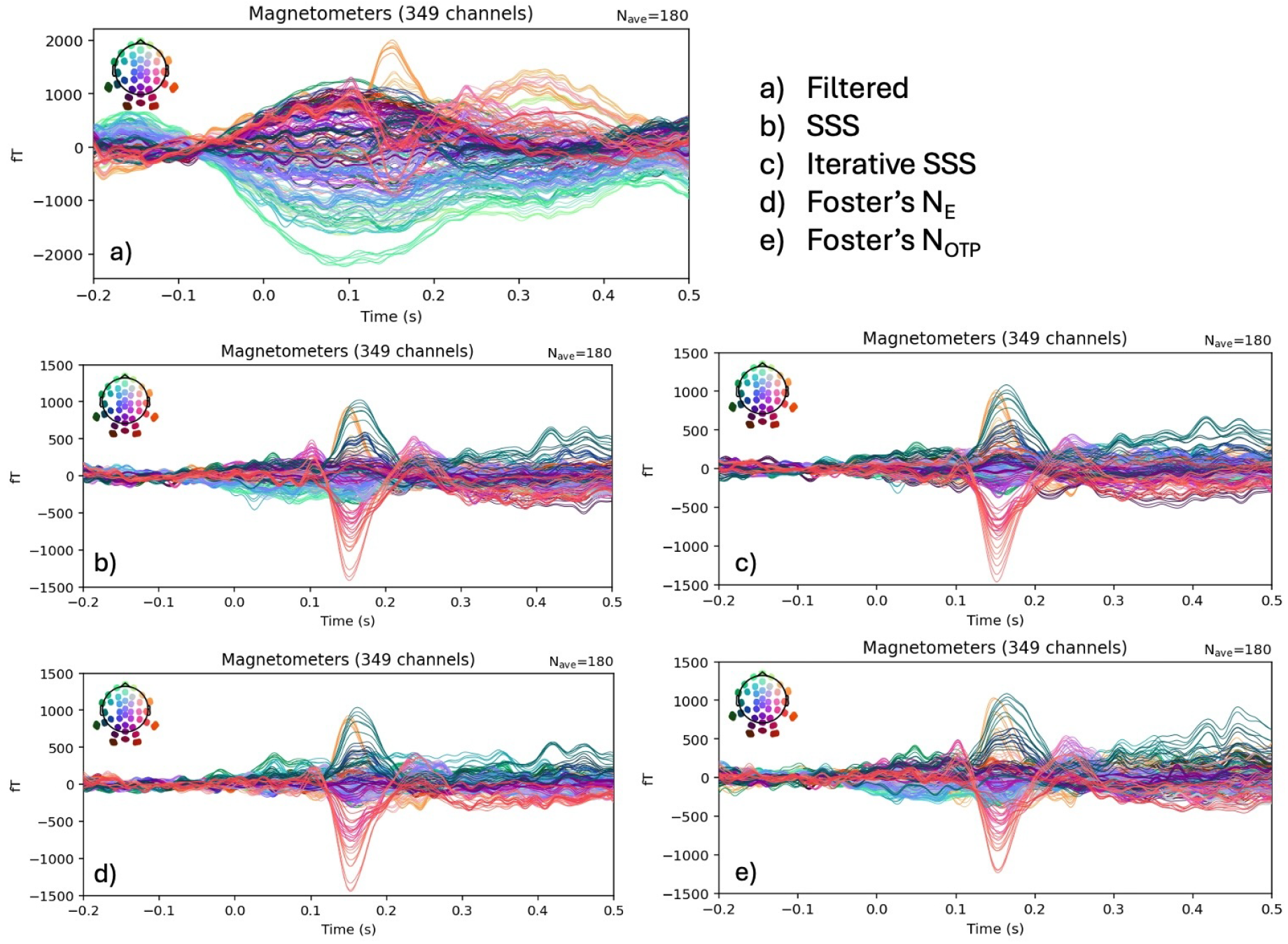
Kernel OPM Audio Evoked butterfly plot for 349 magnetometer channels, baseline corrected, then high-pass-filtered at 0.1 Hz and low-pass filtered at 40 Hz evoked data in Subplot a) is compared to Subplot showing the SSS processed evoked signal, Subplot c) showing the iterative SSS processed evoked signal, Subplot d) Foster’s inverse using empirical sensor noise covariance **N**_E_, and Subplot e) Foster’s inverse using the OTP sensor noise covariance **N**_OTP_.

### 5.4 FieldLine 144-channel OPM Data Set, Stanford University

#### 5.4.1 Empty Room Results

In order to investigate the ability for Foster’s inverse with SSS to combat sensor noise and low frequency fields within the MSR, a two-minute long empty room recording is analyzed with all 144 FieldLine HEDscan OPM sensors on without a human subject in the room. Note that this empty-room recording cannot service as an estimation for the noise covariance profile because the OPM sensors are not in the same position as they would be with a participant inside the helmet. The raw data are then notch filtered at 60 Hz and filtered to allow frequencies between 0.1 Hz and 100 Hz. The frequency spectrum is calculated using MNE-Python for both the filtered raw data and the Foster’s inverse processed raw data for comparison [24].

Compared to the raw empty room frequency spectrum and the other preprocessing methods, Foster’s inverse mitigates the effect of low frequency fields as seen in Figure 9 with a decrease in power for frequencies below 10 Hz. Additionally, the results in Figure 9 shows a reduction in individual sensor noise after Foster’s inverse preprocessing whereas the raw empty room and SSP preprocessed spectrum show a noisy sensor outlying above the others across all frequencies. Finally, Foster’s inverse reduces the variance of noise across all sensors as seen in a reduced thickness of standard deviation in the right column in Figure 9.

**Figure 8.**
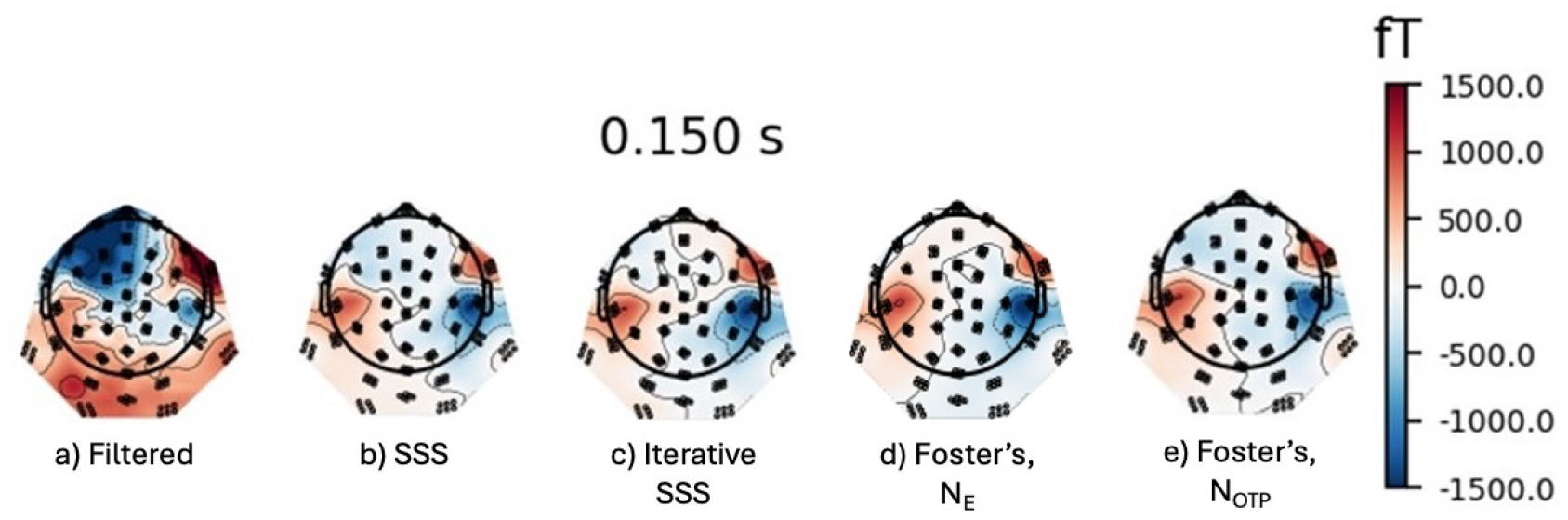
Topographic visualizations of the peak time at 0.15 s of the filtered Kernel OPM Audio Evoked data shown in Panel a) compared to Panel b) showing the SSS-processed evoked auditory response, Panel Iterative SSS with 10 iterations, Panel d) Foster’s inverse using empirical sensor noise covariance **N**_*E*_, and Panel e) Foster’s inverse using the OTP sensor noise covariance **N**_OTP_. Note that only radial-sensing channels are shown in this figure

**Figure 9.**
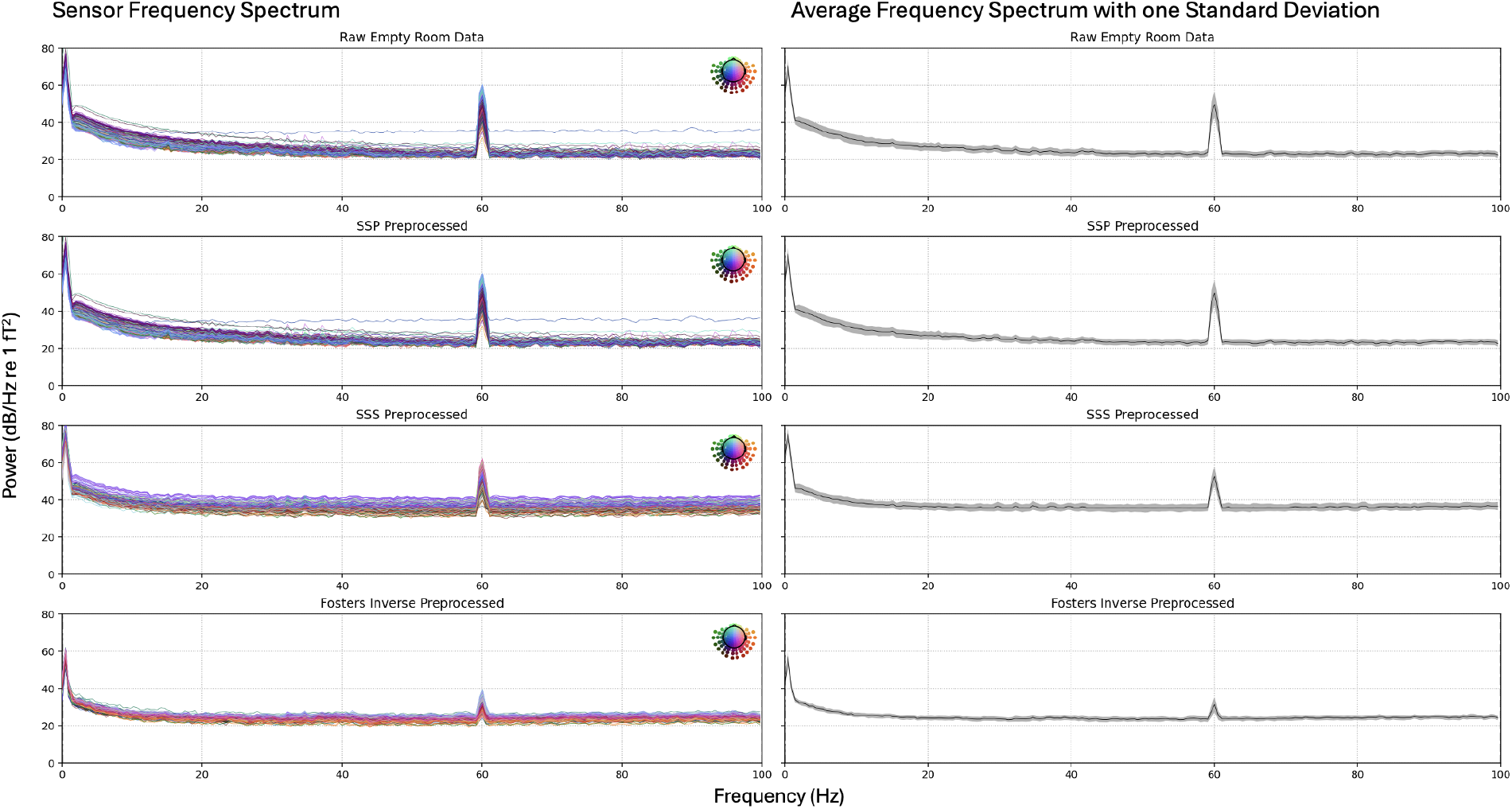
Frequency spectrum from the empty room recorded with the 144 FieldLine OPM sensor system for different preprocessing methods: unfiltered raw empty room data, SSP preprocessed data, SSS preprocessed data, and Foster’s inverse with SSS preprocessed data, from top to bottom. The left column shows the frequency spectrum in Power (dB/Hz re 1 *f T* ^2^) per sensor, and the left column shows the average Power across all sensors at each frequency with ±1 standard deviation in gray highlight.

#### 5.4.2 Noisy Sensor Case Study

To highlight the ability of Foster’s inverse to remove impact from noisy channels, we first focus on one subject where one channel was highly noisy, and is not removed from the data before preprocessing. Before epoching and evoked averaging in MNE-Python, the raw data was minimally high-pass filtered at 0.1 Hz and low-pass filtered at 50 Hz.

The results in Figure 10 compliment the results in Figure 5 and Figure 6, where sensor artifacts and noise are spread through all channels with SSS preprocessing, but Foster’s inverse is able to overcome these artifact by accounting for them in the inverse, reconstructing the internal data without detriments from sensor noise and artifacts. In cases where channels have not been correctly marked bad, many channels are bad, or a sensor became noisier over the course of a recording, Foster’s inverse is able to uncover the brain signals. This case study further exhibits the capabilities of Foster’s inverse; however, it is recommended to remove bad channels before preprocessing methods like SSS to improve the performance and results [36], [37], [10]. Because of this, in further results and subject averages taken with the FieldLine OPM-MEG system, all bad channels have been removed for comparison.

**Figure 10.**
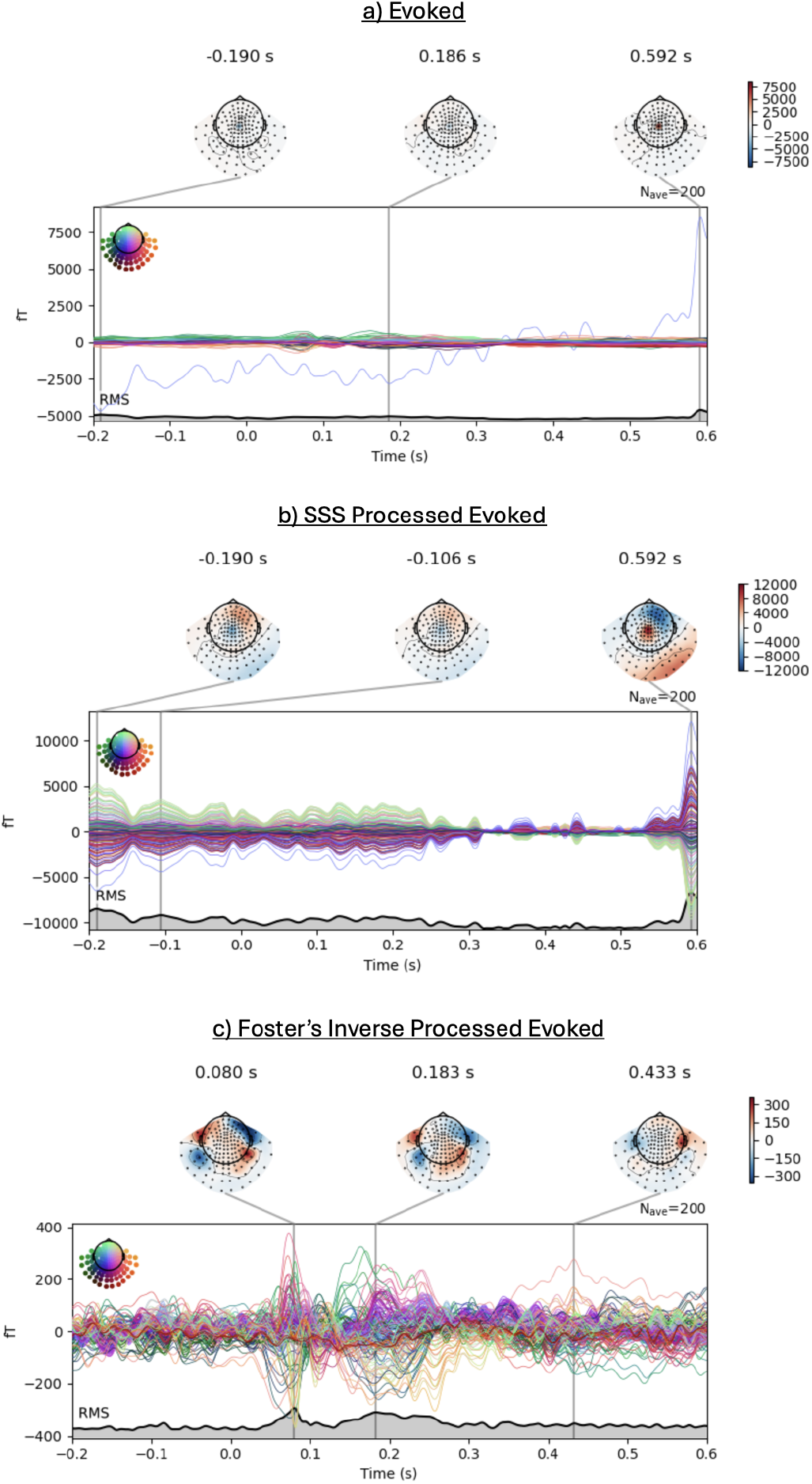
Butterfly Evoked plots and topographic maps of the single-subject case study audio response measured with the FieldLine OPM system. All data was minimally high-pass filtered at 0.1 Hz and low-pass filtered at 50 Hz before calculating evoked responses. Panel a) shows the evoked response, b) shows the evoked response with SSS preprocessing applied, and c) shows the evoked response with Foster’s inverse empirical noise covariance applied.

#### 5.4.3 Evoked and Subject Average Results

Across six subjects, we removed three bad channels. We include comparison with SSP calculated using a baseline period before stimulus onset, as SSP is commonly used in cognitive neuroscience and clinical studies with MEG. We also compare results for two cases of filtering done to the raw data: one is modestly filtered to allow frequencies between 0.5 Hz and 80 Hz with a notch filter at 60 Hz, the other is more heavily filtered between 1 Hz and 50 Hz. First, the average SNR for each preprocessing type is shown in Table 5 with the peak signal calculated between 0.1 s and 0.2 s.

**Table 5:** Six-Subject Average SNR for each preprocessing method.

| Method | 0.5-80 Hz | 1-50 Hz |
| --- | --- | --- |
| Filtered | 1.18 | 2.13 |
| SSP | 1.17 | 3.32 |
| SSS | 1.97 | 4.34 |
| Foster’s Inverse | 4.00 | 6.66 |

Across all subjects, Foster’s inverse with SSS gives the best noise reduction, or highest SNR, as seen in Table 5. The superior noise reduction can be visualized in Figure 11 where the bottom row corresponding to the Foster’s inverse processed data has the clearest evoked topography recovered with automatically detected peaks in both left and right auditory regions as expected, compared to the other methods that are more saturated with noise and drift regardless of how much filtering is applied to the raw data before preprocessing. Additionally, Foster’s inverse provides results that remain stable between the different filters applied to the raw data.

**Figure 11.**
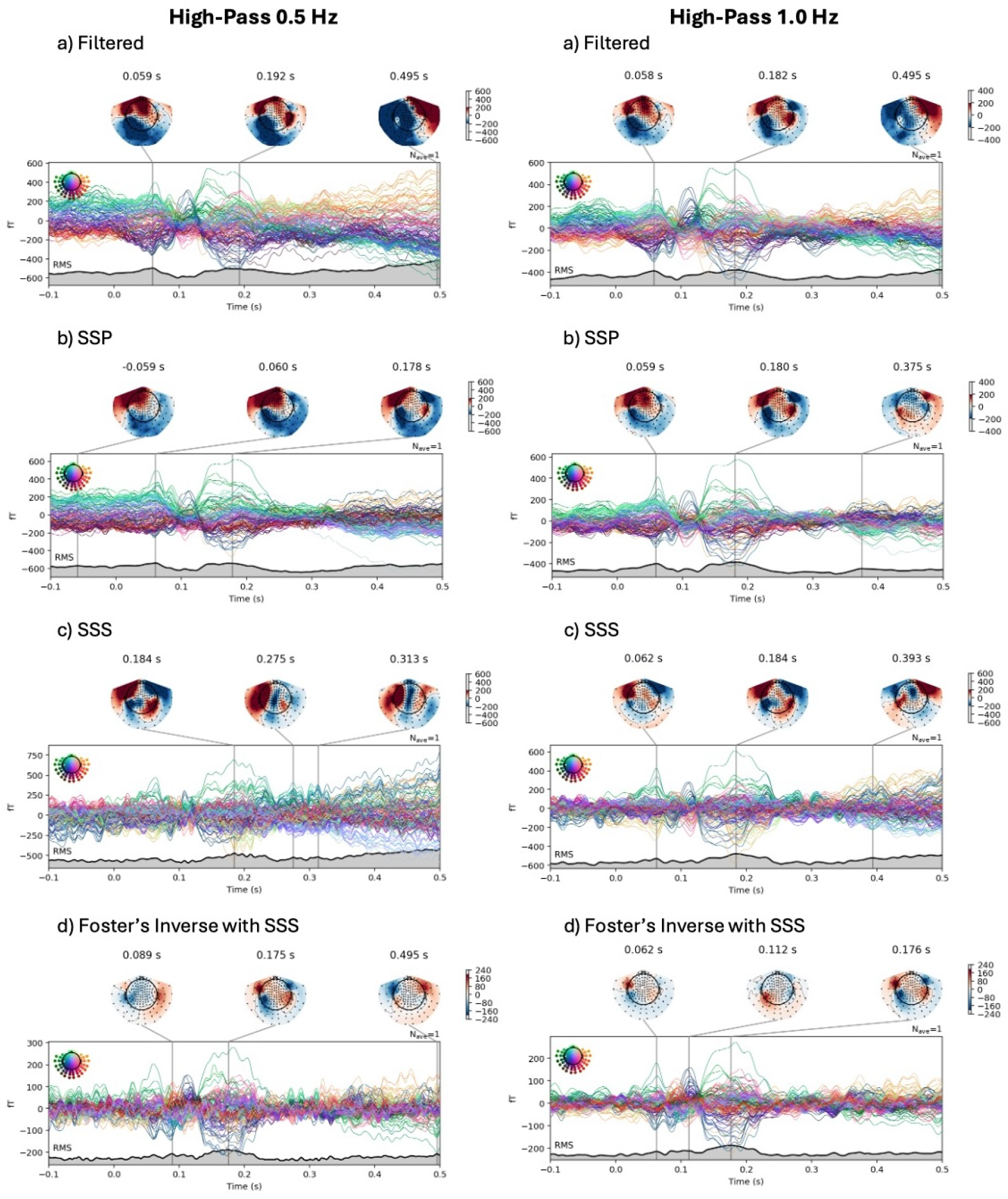
Averaged butterfly evoked plots and topographic maps of six subject audio responses measured with the FieldLine OPM system for two ranges of filtering: the left column contains frequencies from 0.5-80 Hz with a notch filter at 60 Hz, and the right 1-50 Hz. Then, for each frequency range, the raw data is preprocessed with b) SSP, c) SSS, and d) Foster’s inverse with SSS before evoked averaging.

**Figure 12.**
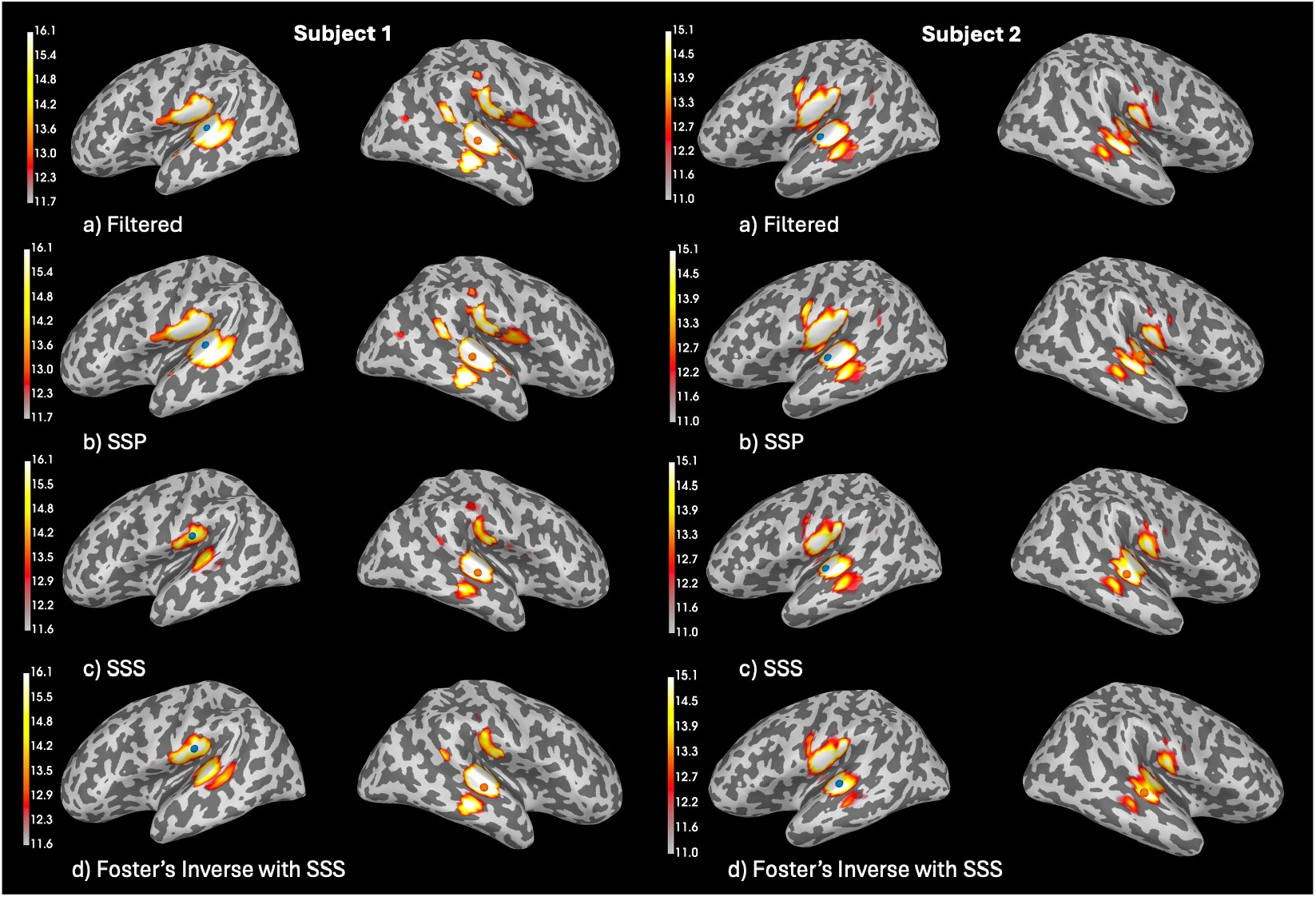
Source localization results for Subject 1 in the left column at peak time 0.107 s (left) and Subject 2 in the right column at peak time 0.175 s (right) with dSPM inverse solution and free orientation dipoles. Before forward and inverse modeling, the raw data is preprocessed using one of the four preprocessing methods show in row a) Filtered with a high-pass at 0.5 Hz, low-pass at 80 Hz, and a notch at 60 Hz, row b) SSP, row c) SSS, and row d) Foster’s inverse with SSS.

#### 5.4.4 Source Localization Results

Here, we present the source localization results for three subjects with individual structural MRIs. The explained variance (EV %) and minimum/maximum GOF (%) for the best dipole fit are shown in Table 6. We provide visuals of the source localization at the peak time in MRI space for Subject 1 and Subject 2 in Figure 14.

**Table 6:** Source Localization Metrics for three subject audio responses measured using FieldLine OPM sensors.

| Subject | method | EV (%) | Min/Max GOF (%) |
| --- | --- | --- | --- |
| S1 | Filtered | 90.1 | 12.1/27.4 |
|  | SSP | 90.1 | 12.0/28.0 |
|  | SSS | 91.1 | 16.5/34.7 |
|  | Foster’s Inverse | 91.9 | 17.4/33.8 |
| S2 | Filtered | 85.7 | 12.4/31.0 |
|  | SSP | 85.6 | 10.6/31.1 |
|  | SSS | 88.3 | 14.5/44.6 |
|  | Foster’s Inverse | 88.0 | 19.3/37.0 |
| S3 | Filtered | 86.1 | 13.1/52.0 |
|  | SSP | 86.5 | 12.8/52.3 |
|  | SSS | 88.9 | 18.7/60.2 |
|  | Foster’s Inverse | 89.1 | 18.9/57.6 |
| **Average** | Filtered | 85.6 | 11.6/38.8 |
|  | SSP | 85.7 | 11.2/39.0 |
|  | SSS | 89.8 | 16.6/46.5 |
|  | Foster’s Inverse | 89.7 | 18.5/42.8 |

The GOF percentages reported in Table 6 are relatively low when compared to the Phantom data in Table 2 and 3 because the dipole fitting function finds the location and activity of a singular dipole source at each time point, which is ideal for Phantom recordings where each dipole is activated in sequence. But, as seen in Figure 14, auditory activity occurs bilaterally, so the activity is from multiple sources simultaneously. Thus, the GOF can be interpreted as the best fit for one dipole that best explains the majority of the activity at that one point in time. On average, Foster’s inverse performs better than high-pass filtering and SSP, and performs as well as SSS in EV% and GOF %, but offers better source localization with an increase in minimum GOF % as seen in Table 14 and with smaller activation probability areas as seen in Figure 14.

#### 5.4.5 Foster’s Inverse with mSSS

Here, we demonstrate how Foster’s inverse can be used with the mSSS method instead of the SSS method for calculating the spatial basis expansion and the corresponding multipole moments [27]. First, we replicate the process for calculating averaged evoked responses, but with the same subset of three subjects with individual MRIs. Next, we calculate the explained variance both the minimum and maximum GOF across the 102 source localized dipoles in Table 7, with corresponding visualizations at the peak time.

**Table 7:** Source Localization Metrics for three subject audio responses measured using FieldLine OPM sensors with three different SSS-based preprocessing techniques.

| Subject | method | EV (%) | Min/Max GOF (%) |
| --- | --- | --- | --- |
| S1 | SSS | 92.1 | 16.5/34.7 |
|  | Foster's Inverse | 91.9 | 17.4/33.8 |
|  | Foster's Inverse with mSSS | 91.3 | 25.3/46.3 |
| S2 | SSS | 88.3 | 14.5/44.6 |
|  | Foster's Inverse | 88.0 | 19.3/37.0 |
|  | Foster's Inverse with mSSS | 89.5 | 20.1/41.6 |
| S3 | SSS | 88.9 | 18.7/60.2 |
|  | Foster's Inverse | 89.1 | 18.9/57.6 |
|  | Foster's Inverse with mSSS | 87.2 | 24.1/63.0 |
| **Average** | SSS | 89.8 | 16.6/46.5 |
|  | Foster's Inverse | 89.7 | 18.5/42.8 |
|  | Foster's Inverse with mSSS | 89.4 | 23.2/50.3 |

In figure 13, the subject-average amplitude of the peak signal is higher for the mSSS case than the SSS, potentially due to the mSSS basis providing a better span of the brain-space and capturing all of the internal signals that may have been excluded with only a single-origin expansion [27]. Foster’s inverse with mSSS increases the goodness of fit of the source localization on average as seen in Table 7, performs similarly in explained variance, and uncovers the same source localization results with reduced area as seen in Figure 14, complimenting the findings of increased goodness of fit. These results indicate that Foster’s inverse with mSSS can further improve source localization results with on-scalp MEG systems like OPMs, and shows that Foster’s inverse as a data preprocessing method is highly adaptable to different methods of obtaining the spatial expansions of the brain’s magnetic signals.

**Figure 13.**
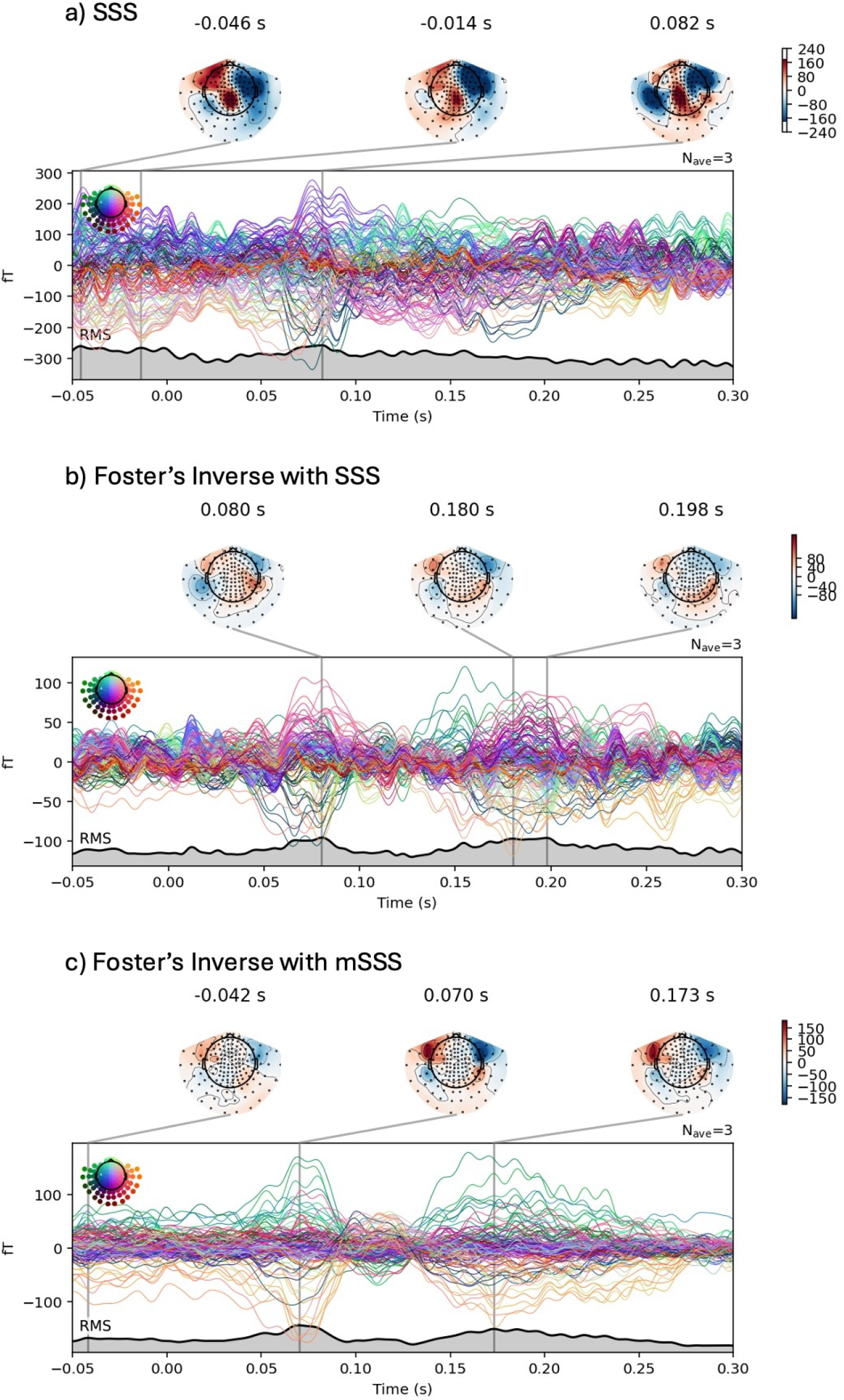
Butterfly Evoked plots and topographic maps three-subject grand averaged response measured with the FieldLine OPM system. All data was minimally high-pass filtered at 0.5 Hz and low-pass filtered at 50 Hz before calculating evoked responses. The Panel a) shows the evoked response with SSS preprocessing applied, Panel b) shows the evoked response with Foster’s inverse with SSS, and Panel c) shows Foster’s inverse with mSSS.

**Figure 14.**
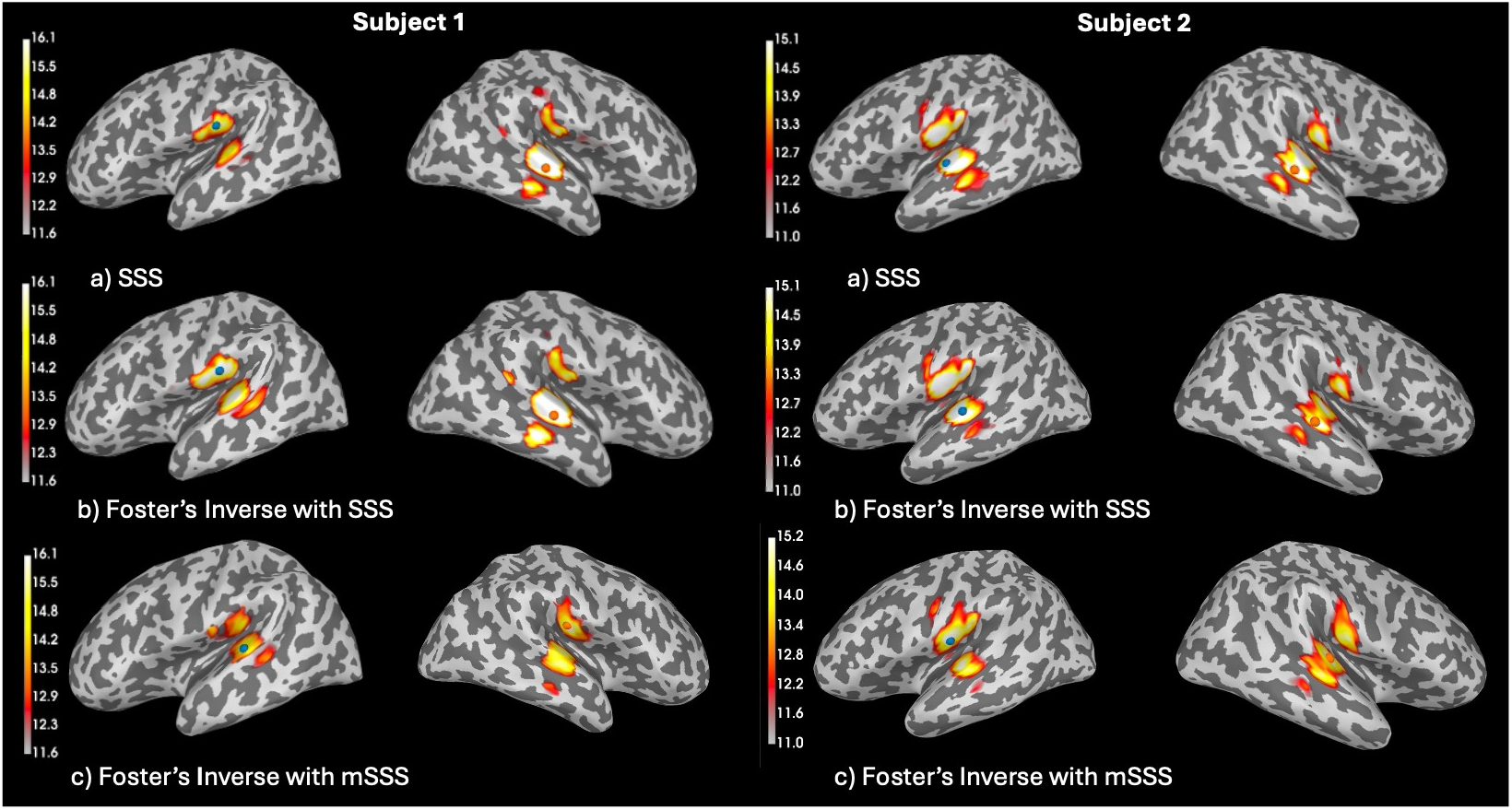
Source localization results for Subject 1 in the left column at peak time 0.107 s (left) and Subject 2 in the right column at peak time 0.175 s (right) with dSPM inverse solution and free orientation dipoles. Before forward and inverse modeling, the raw data is filtered with a notch at 60 Hz, high-pass at 0.5 Hz and a low-pass at 80 Hz, then preprocessed with SSS in row a), Foster’s inverse with SSS in row b), and Foster’s inverse with mSSS in row c).

## 6 Comments and Conclusions

In this paper, we explored the application of Foster’s matrix inverse method to cryogenic MEG and OPM-MEG data. We tested whether this method leads to an improved reconstruction of the magnetic fields generated by brain activity, free from exterior signals and sensor artifacts. Overall, we find that Foster’s inverse with SSS substantially improves SNR and localisation accuracy as compared to other commonly used methods. Foster’s inverse therefore holds significant promise as a preprocessing step for improving data quality, and we have implemented this method utilizing existing MNE-Python functionality to allow others to easily adopt this approach.

Through noisy current dipole simulations, we found that Foster’s inverse with SSS substantially improves the SNR of the reconstructed data by a factor of 10 as seen in Table 1 when the noise covariance matrix is calculated from the Gaussian noise generated in the simulation. We obtained similar results when testing Foster’s inverse on data from a Phantom head containing 32 dipoles measured using a SQUID MEG system. Foster’s inverse provides improved dipole localization and GOF percentages across different recording days and dipole activation strengths, and the method can overcome significant sensor artifacts where traditional SSS fails. The use of Foster’s inverse is particularity important in cases where artifacts or bad channels are missed through artifact and bad channel detection methods or protocols, protecting against undetected spreading of noise and allowing for robust reconstructions of the internal brain signals.

Finally, we tested Foster’s inverse with two different OPM-MEG systems: the 432-channel Kernel Flux OPM system at the University of Washington, and the 144-channel FieldLine HEDscan OPM system at Stanford University. Foster’s inverse with noise covariance calculated using the empirical method offers improved noise reduction, particularly with the presence of an empty-room recording used to obtain the calculation. With the FieldLine system, we first examined a set of empty room recordings which were then preprocessed using SSP, SSS, and Foster’s inverse. Foster’s inverse reduces the low-frequency fields present in the MSR, as well as reducing the amplitude and spread of the noise across all sensors, highlighting its ability to combat the types of noise specific to OPM-MEG sensors. As seen in the SQUID MEG Phantom results, Foster’s inverse is robust against OPM-MEG sensor artifacts as well, whereas SSS suffers from the spreading of sensor artifacts across channels. Across six subjects listening to the same audio tone stimulus, Foster’s inverse reconstructs brain activity and removes noise, recovering the expected topographic maps for auditory cortex activation.

For three of these subjects with structural MRIs, forward and inverse results were calculated after each of the four preprocessing methods were applied to the data (filtering between 0.5 and 80 Hz with a notch at 60 Hz, SSP, SSS, and Foster’s inverse). Inverse modeling was done using dSPM in MNE-Python with free dipole orientations. Here, Foster’s inverse offers improvements in SNR, source localization, and goodness of fit of the localized dipole source. For the FieldLine OPM system, the goodness of fit results are further improved when using Foster’s inverse in combination with the mSSS method for determining the spatial basis expansion components.

Foster’s inverse relies on an accurately measured covariance matrix **N** that only contains sensor noise. We discussed different methods for calculating the sensor noise covariance, including the Empirical covariance method implemented in the MNE-Python package, as well as other novel possibilities including the use of OTP to isolate random sensor noise from the measured data [24], [23]. When it comes to selecting a method for obtaining the spatial basis components and sensor noise covariance, the results in this study offer some suggestions. First, using methods for obtaining the spatial components that have been optimized for different types of MEG systems, such as SSS for cryogenic-MEG and mSSS for OPM-MEG, offers improved results. Second, the type of MEG system and available data influences the choice for the noise covariance calculation method. In general, for MEG and OPM-MEG recordings with both correlated and spatially uncorrelated sensor noise, the empirical method performs best when used in Foster’s inverse. Due to the importance of the noise covariance profile on the functionality of Foster’s inverse with SSS, methods for obtaining the profile specifically for OPM and other on-scalp systems will need to be further investigated as the technology becomes more prevalent. Here, we demonstrated that Foster’s inverse is highly adaptable and robust to multiple methods for calculating noise covariance and the basis expansion of the magnetic fields.

Overall, Foster’s inverse with SSS is a novel, robust, and powerful matrix inversion method for combating sensor noise, resulting in improved source localization over other MEG preprocessing methods. As the field of cognitive neuroscience increasingly utilizes OPM-MEG technology, Foster’s inverse can be used to mitigate OPM sensitivity to low-frequency fields and variable noise floors. This allows the sensors to offer improved source localization through increased sensitivity to higher-order spatial components of the neuronal magnetic fields which were not previously captured in such details by the off-scalp cryogenic MEG sensor systems. We have shown the functionality and power of Foster’s inverse to reduce noise, remove the need for aggressive high-pass filtering, and ultimately uncover the higher spatial frequencies needed for better source localization accuracy with cryogenic MEG, OPM-MEG, and other on-scalp systems.

**7 Appendix**

### 7.1 Multipole Moment Formula

The coefficients *α*_*lm*_ and *β*_*lm*_ in Equation 3 denote the multiple moments and can be written in a lead-field like representation using the vector spherical harmonic function **X**_*lm*_(*θ, φ*)

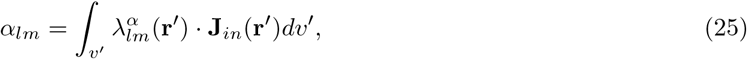

where **J**_*in*_(**r**^′^) is the internal current distribution and *λ*^*α*^ (**r**^′^) represent the analogy to the conventional lead fields. Fully expanded, we have

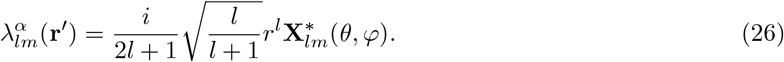

**X**_*lm*_(*θ, φ*) can be written using the spherical harmonic function *Y*_*lm*_(*θ, φ*) as [34]

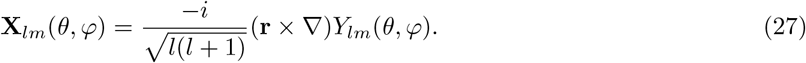

Note that the current **J** given from Maxwell’s equations as the curl of the magnetic field must be zero given the VSH expansion of **B**, meaning the sensors must be located in a source-free zone outside of the head.

### 7.2 Analogy to the Linear Wiener Estimate in Noise-Normalized MNE Methods

As mentioned here and extensively studied, the inverse problem of mapping collected MEG (and EEG) data into the location and time of brain currents creating the signal has, in principle, infinite solutions. Thus, many have developed methods to constrain the most likely solutions to the inverse problem using what we know about biology, brain structure, and the underlying physics of the signals. One large category of these solutions are Minimum-Norm Estimates (MNEs), which on a simple level, solve the noiseless equation ***y*** = **A*x*** with the MNE estimate for the solution 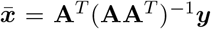, where ***y*** is the measurement vector, **A** is some mixing matrix showing the mapping of the currents onto the sensors, and ***x*** are the unknown current sources [22]. MNE methods can be improved using noise-normalization techniques, such as dynamic statistical parameter mapping (dSPM) [7] and standardized low resolution brain electromagnetic tomography (sLORETA) [29], which weight the MNE solution to combat the fact that MNE methods are biased to locate sources superficially on the cortex that in reality originated from deeper activity [22]. Both methods begin with the same data vector ***y*** = **A*x*** + ***η*** as discussed previously in the Foster’s inverse section. Estimates for the brain current sources 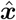 are obtained using the Linear Wiener estimate **W** [44] where

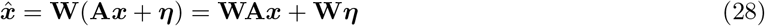

The Wierner estimator is written in terms of the spatial covariance of the dipole strengths **R** = ⟨***x***(*t*)***x***(*t*)⟩and the covariance of the sensor noise vector **N** = ⟨***η***(*t*)***η***(*t*)⟩ as [7],[22]:

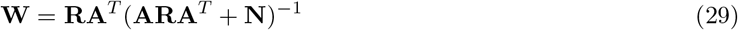

Comparing this equation to Equation 17, we can see that this form of **W** shares similar mathematical composition with **B** from Foster’s inverse, where the covariance of the multipole moments matrix **X** is analogous to the spatial covariance of the dipole strength matrix **R**, and the SSS matrix **S** is analogous to the forward matrix operator **A** [7], or mixing matrix [22].

The full Foster’s inverse solution for the current multipole moments 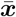 can written as follows by plugging in **B** and ***b*** in Equation 16:

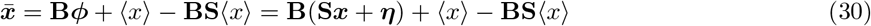

Where the first term **B**(**S*x*** + ***η***) matches the Wierner estimate for the brain currents 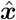 in Equation 28. The complete Foster’s solution in Equation 30 contains extra terms pertaining to the average multipole moments found during SSS, which are further weighted by the SSS matrix itself in the term ***b***.

### 7.3 The Case of Minimum Information

The aforementioned presentation of Foster’s inverse in SSS requires knowledge about the noise covariance matrix **N** in order for the method to function as stated. In practice, knowing the exact profile of the MEG sensor noise can be quite difficult. In the minimum information case, the full profile of the noise is unknown, but the covariance matrix can be written as 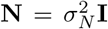 where 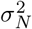 is the noise power and **I** is an identity matrix with the same dimension as matrix **N** [13]. Similarly, the covariance of multipole moments can be written as 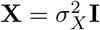 where 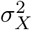 is the signal power. Using these variables, we can rewrite Equation 17 as

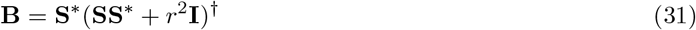

where 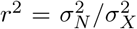 is the noise-to-signal power ratio [13]. Specifically in the context of SSS, 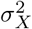 corresponds to the SNR of the multipole moments. Note that the minimum-information inverse operator **B** has the same structure as the regularized Tikhonov operator **T** which gives solutions to noisy linear equations, given by

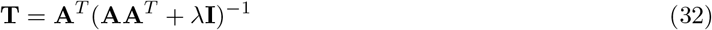

where *λ* is some positive regularization parameter [22]. By comparing **B** with **T**, we see that Foster’s inverse can offer one possibility to the value of *λ* using the ratio 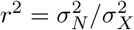, which acts as a regularization parameter in the minimum-information case of Foster’s inverse. In fact, choosing 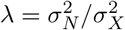 is the unbiased, linear, and minimum mean square error estimator of ***x*** [22]. Using Foster’s inverse in this fashion means that the maximum amplification factor from any one large instance of sensor noise impacts the solutions by a factor of 1*/*(2*r*) regardless of the amplitudes of the SSS basis expansion [13]. Estimating or calculating the noise-to-signal power ratio may be simpler than finding the full noise covariance profile, especially for on-scalp MEG systems that have not been widely characterized.

## 8 Ethics Statement

All data were collected in accordance with the given university’s institutional review board (IRB) human subjects testing guidelines, and informed consent was obtained from all participants for being included in the study.

## 9 Data and Code Availability

The Python implementation Foster’s inverse for SSS is available online [25], and is designed to compliment the implementation of SSS in MNE-Python [24], where an integration of the two is in progress. The Foster’s implementation prompts users to choose the MNE-Python empirical method or the OTP method for calculation of the noise covariance [24], [10], [23]. Single-subject Kernel Flux data is available online [26]. Phantom cryogenic MEG and FieldLine OPM-MEG data are available upon request, based on the need for a formal data sharing agreement.

## 10 Author Contributions

**Alexandria McPherson**: conceptualization, methodology, software, validation, formal analysis, investigation, resources, data simulation, data collection, data processing, visualization, writing (original draft, review, and editing), project administration. **Sepp Sanchirico**: software, validation, formal analysis, data simulation, visualization. **Albert Xu**: software, validation, formal analysis, data simulation, visualization. **Eric Larson**: conceptualization, methodology, software, validation, resources, writing (review and editing). **Milena Kaestner**: data collection, writing (review and editing) **William Turner**: software, writing (review and editing) **Laura Gwilliams**: conceptualization, methodology, resources, visualization, writing (review and editing), supervision, funding acquisition. **Samu Taulu**: conceptualization, methodology, resources, visualization, writing (review and editing), supervision, funding acquisition. *Correspondence* should be sent to A.M.

## 11 Funding

This work was supported by grants R21-EB033577-01 and R01-NS104585-01A1 from the National Institutes of Health. S. Taulu’s work is also funded in part by the Bezos Family Foundation and the R. B. and Ruth H. Dunn Charitable Foundation. LG was funded by the Esther A. & Joseph Klingenstein Fund; Whitehall Foundation 2024-08-043; BRAIN Foundation A-0741551370.

## 12 Declaration of Competing Interest

The authors have no conflicts of interest relevant to this work.

## 13 Acknowledgments

We acknowledge the Koret Human Neurosciences Community Laboratory.

## 14 Supplementary Materials

## Notes

https://zenodo.org/records/21079671

### Competing Interest Statement

The authors have declared no competing interest.

## References

[1] Orang Alem et al. “An integrated full-head OPM-MEG system based on 128 zero-field sensors”. In: Frontiers in Neuroscience 17 (June 2023), p. 1190310. ISSN: 1662-4548. DOI: 10.3389/fnins.2023.1190310. URL: https://pmc.ncbi.nlm.nih.gov/articles/PMC10303922/ (visited on 02/18/2026).

[2] Richard Aveyard, Joe Lyons, and Alexander Robert Wade. York OPM Registration Code (YORC). Dec. 2024. DOI: 10.5281/zenodo.20119082. URL: https://zenodo.org/records/20119082 (visited on 05/11/2026).

[3] Elena Boto et al. “Moving magnetoencephalography towards real-world applications with a wearable system”. en. In: Nature 555.7698 (Mar. 2018), pp. 657–661. ISSN: 1476-4687. DOI: 10.1038/nature26147. URL: https://www.nature.com/articles/nature26147 (visited on 07/22/2025).

[4] Matthew J. Brookes et al. “Magnetoencephalography with optically pumped magnetometers (OPM-EG): the next generation of functional neuroimaging”. en. In: Trends in Neurosciences 45.8 (Aug. 2022), pp. 621–634. ISSN: 01662236. DOI: 10.1016/j.tins.2022.05.008. URL: https://linkinghub.elsevier.com/retrieve/pii/S0166223622001023 (visited on 09/26/2024).

[5] Alain de Cheveigné and Jonathan Z. Simon. “Sensor noise suppression”. In: Journal of neuroscience methods 168.1 (Feb. 2008), pp. 195–202. ISSN: 0165-0270. DOI: 10.1016/j.jneumeth.2007.09.012. URL: https://www.ncbi.nlm.nih.gov/pmc/articles/PMC2253211/ (visited on 07/22/2025).

[6] Maggie Clarke et al. Human Brain Mapping — Neuroimaging Journal — Wiley Online Library. Mar. 2022. URL: https://onlinelibrary.wiley.com/doi/10.1002/hbm.25871?msockid=3fc1acc8af5d6caf258abe2faee76d1a (visited on 07/21/2025).

[7] Anders M Dale et al. “Mapping: Combining fMRI and MEG for High-Resolution Imaging of Cortical Activity”. en. In: (2000).

[8] Joram van Driel, Christian N. L. Olivers, and Johannes J. Fahrenfort. “High-pass filtering artifacts in multivariate classification of neural time series data”. In: Journal of Neuroscience Methods 352 (Mar. 2021), p. 109080. ISSN: 0165-0270. DOI: 10.1016/j.jneumeth.2021.109080. URL: https://www.sciencedirect.com/science/article/pii/S0165027021000157 (visited on 02/18/2026).

[9] ELEKTA. Elekta Neuromag Technical Manual, Revision G. Sept. 2005. URL: https://www.manualslib.com/manual/1482268/Elekta-Neuromag.html (visited on 09/12/2025).

[10] Denis A. Engemann and Alexandre Gramfort. “Automated model selection in covariance estimation and spatial whitening of MEG and EEG signals”. In: NeuroImage 108 (Mar. 2015), pp. 328–342. ISSN: 1053-8119. DOI: 10.1016/j.neuroimage.2014.12.040. URL: https://www.sciencedirect.com/science/article/pii/S1053811914010325 (visited on 08/14/2025).

[11] Bruce Fischl. “FreeSurfer”. In: NeuroImage. 20 YEARS OF fMRI 62.2 (Aug. 2012), pp. 774–781. ISSN: 1053-8119. DOI: 10.1016/j.neuroimage.2012.01.021. URL: https://www.sciencedirect.com/science/article/pii/S1053811912000389 (visited on 06/18/2026).

[12] K. B. Fisher et al. “Wiener reconstruction of density, velocity and potential fields from all-sky galaxy redshift surveys”. In: Monthly Notices of the Royal Astronomical Society 272.4 (Feb. 1995), pp. 885–908. ISSN: 0035-8711. DOI: 10.1093/mnras/272.4.885. URL: https://doi.org/10.1093/mnras/272.4.885 (visited on 10/23/2025).

[13] Manus Foster. “An Application of the Wiener-Kolmogorov Smoothing Theory to Matrix Inversion”. In: Journal of the Society for Industrial and Applied Mathematics 9.3 (Sept. 1961), pp. 387–392. ISSN: 0368-4245. DOI: 10.1137/0109031. URL: https://epubs.siam.org/doi/10.1137/0109031 (visited on 02/21/2025).

[14] L. Gwilliams, G. A. Lewis, and A. Marantz. “Functional characterisation of letter-specific responses in time, space and current polarity using magnetoencephalography”. In: NeuroImage 132 (May 2016), pp. 320–333. ISSN: 1053-8119. DOI: 10.1016/j.neuroimage.2016.02.057. URL: https://www.sciencedirect.com/science/article/pii/S105381191600166X (visited on 04/17/2026).

[15] Laura Gwilliams et al. “Hierarchical dynamic coding coordinates speech comprehension in the human brain”. In: Proceedings of the National Academy of Sciences 122.42 (Oct. 2025), e2422097122. DOI: 10.1073/pnas.2422097122. URL: https://www.pnas.org/doi/10.1073/pnas.2422097122 (visited on 04/06/2026).

[16] Matti Hämäläinen et al. “Magnetoencephalography—theory, instrumentation, and applications to non-invasive studies of the working human brain”. en. In: Reviews of Modern Physics 65.2 (Apr. 1993), pp. 413–497. ISSN: 0034-6861, 1539-0756. DOI: 10.1103/RevModPhys.65.413. URL: https://link.aps.org/doi/10.1103/RevModPhys.65.413 (visited on 09/23/2024).

[17] Liisa Helle et al. “Extended Signal-Space Separation Method for Improved Interference Suppression in MEG”. In: IEEE Transactions on Biomedical Engineering 68.7 (July 2021), pp. 2211–2221. ISSN: 1558-2531. DOI: 10.1109/TBME.2020.3040373. URL: https://ieeexplore.ieee.org/document/9268467 (visited on 04/07/2026).

[18] Niall Holmes et al. “An Iterative Implementation of the Signal Space Separation Method for Magnetoencephalography Systems with Low Channel Counts”. en. In: Sensors 23.14 (Jan. 2023). Number: 14, p. 6537. ISSN: 1424-8220. DOI: 10.3390/s23146537. URL: https://www.mdpi.com/1424-8220/23/14/6537 (visited on 12/13/2023).

[19] Niall Holmes et al. “Wearable magnetoencephalography in a lightly shielded environment”. In: IEEE Transactions on Biomedical Engineering (2024). Conference Name: IEEE Transactions on Biomedical Engineering, pp. 1–10. ISSN: 1558-2531. DOI: 10.1109/TBME.2024.3465654. URL: https://ieeexplore.ieee.org/document/10685146 (visited on 09/26/2024).

[20] Mingxiong Huang, Mårten Risling, and Dewleen G. Baker. “The role of biomarkers and MEG-based imaging markers in the diagnosis of post-traumatic stress disorder and blast-induced mild traumatic brain injury”. In: Psychoneuroendocrinology 63 (Jan. 2016), pp. 398–409. ISSN: 0306-4530. DOI: 10.1016/j.psyneuen.2015.02.008. URL: https://www.sciencedirect.com/science/article/pii/S030645301500058X (visited on 07/21/2025).

[21] Joonas Iivanainen. “Spatial-jitter model for magnetoencephalography sensor arrays”. In: IEEE Transactions on Medical Imaging PP (Apr. 2026), pp. 1–1. DOI: 10.1109/TMI.2026.3687982.

[22] Risto J. Ilmoniemi and Jukka Sarvas. Brain Signals: Physics and Mathematics of MEG and EEG. en. Google-Books-ID: wZiWDwAAQBAJ. MIT Press, May 2019. ISBN: 978-0-262-35282-6.

[23] Eric Larson and Samu Taulu. “Reducing Sensor Noise in MEG and EEG Recordings Using Oversampled Temporal Projection”. In: IEEE Transactions on Biomedical Engineering 65.5 (May 2018), pp. 1002–1013. ISSN: 1558-2531. DOI: 10.1109/TBME.2017.2734641. URL: https://ieeexplore.ieee.org/document/7997929 (visited on 07/23/2025).

[24] Eric Larson et al. MNE-Python. Dec. 2024. DOI: 10.5281/zenodo.14519545. URL: https://zenodo.org/records/14519545 (visited on 02/21/2025).

[25] Alexandria McPherson. Fosters Inverse for SSS. Apr. 2026. DOI: 10.5281/zenodo.19463038. URL: https://zenodo.org/records/19463038 (visited on 04/07/2026).

[26] Alexandria McPherson et al. “2025-Refined-SSS-Methods-On-Scalp-MEG”. en. In: (Feb. 2025). URL:https://osf.io/teygz/ (visited on 05/12/2025).

[27] Alexandria N McPherson et al. “Refined signal space separation methods for on-scalp MEG systems”. en. In: Physics in Medicine & Biology (2025). ISSN: 0031-9155. DOI: 10.1088/1361-6560/ade6ba. URL: http://iopscience.iop.org/article/10.1088/1361-6560/ade6ba (visited on 06/25/2025).

[28] Hiroatsu Murakami et al. “Correlating magnetoencephalography to stereo-electroencephalography in patients undergoing epilepsy surgery”. In: Brain 139.11 (Nov. 2016), pp. 2935–2947. ISSN: 0006-8950. DOI: 10.1093/brain/aww215. URL: https://doi.org/10.1093/brain/aww215 (visited on 07/21/2025).

[29] R. D. Pascual-Marqui. “Standardized low-resolution brain electromagnetic tomography (sLORETA): technical details”. eng. In: Methods and Findings in Experimental and Clinical Pharmacology 24 Suppl D (2002), pp. 5–12. ISSN: 0379-0355.

[30] Mangor Pedersen, David F. Abbott, and Graeme D. Jackson. “Wearable OPM-MEG: A changing landscape for epilepsy”. en. In: Epilepsia 63.11 (2022). eprint: https://onlinelibrary.wiley.com/doi/pdf/10.1111/epi.17368, pp. 2745–2753. ISSN: 1528-1167. DOI: 10.1111/epi.17368. URL: https://onlinelibrary.wiley.com/doi/abs/10.1111/epi.17368 (visited on 10/23/2025).

[31] Jonathan Peirce et al. “PsychoPy2: Experiments in behavior made easy”. en. In: Behavior Research Methods 51.1 (Feb. 2019), pp. 195–203. ISSN: 1554-3528. DOI: 10.3758/s13428-018-01193-y. URL: https://doi.org/10.3758/s13428-018-01193-y (visited on 04/06/2026).

[32] Ethan J. Pratt et al. “Kernel Flux: Optical and Quantum Sensing and Precision Metrology 2021”. In: Optical and Quantum Sensing and Precision Metrology. Proceedings of SPIE - The International Society for Optical Engineering (2021). Ed. by Selim M. Shahriar and Jacob Scheuer. DOI: 10.1117/12.2581794. URL: http://www.scopus.com/inward/record.url?scp=85106763749–partnerID=8YFLogxK (visited on 10/15/2024).

[33] J Sarvas. “Basic mathematical and electromagnetic concepts of the biomagnetic inverse problem”. en. In: Physics in Medicine and Biology 32.1 (Jan. 1987), pp. 11–22. ISSN: 0031-9155, 1361-6560. DOI: 10.1088/0031-9155/32/1/004. URL: https://iopscience.iop.org/article/10.1088/0031-9155/32/1/004 (visited on 01/18/2024).

[34] Samu Taulu and Matti Kajola. “Presentation of electromagnetic multichannel data: The signal space separation method”. In: Journal of Applied Physics 97.12 (June 2005), p. 124905. ISSN: 0021-8979. DOI: 10.1063/1.1935742. URL: https://doi.org/10.1063/1.1935742 (visited on 12/13/2023).

[35] Samu Taulu and Juha Simola. “Spatiotemporal signal space separation method for rejecting nearby interference in MEG measurements”. en. In: Physics in Medicine & Biology 51.7 (Mar. 2006), p. 1759. ISSN: 0031-9155. DOI: 10.1088/0031-9155/51/7/008. URL: https://dx.doi.org/10.1088/0031-9155/51/7/008 (visited on 12/13/2023).

[36] Samu Taulu, Juha Simola, and Matti Kajola. “Applications of the signal space separation method”. In: IEEE Transactions on Signal Processing 53.9 (Sept. 2005). Conference Name: IEEE Transactions on Signal Processing, pp. 3359–3372. ISSN: 1941-0476. DOI: 10.1109/TSP.2005.853302. URL: https://ieeexplore.ieee.org/document/1495874 (visited on 12/13/2023).

[37] Samu Taulu et al. “Novel Noise Reduction Methods”. In: Magnetoencephalography: From Signals to Dynamic Cortical Networks. Journal Abbreviation: Magnetoencephalography: From Signals to Dynamic Cortical Networks. July 2014, pp. 35–71. ISBN: 978-3-642-33044-5. DOI: 10.1007/978-3-642-33045-2_2.

[38] Tim M Tierney et al. Adaptive multipole models of OPM data. en. preprint. Neuroscience, Sept. 2023. DOI: 10.1101/2023.09.11.557150. URL: http://biorxiv.org/lookup/doi/10.1101/2023.09.11.557150 (visited on 12/13/2023).

[39] Tim M. Tierney et al. “Modelling optically pumped magnetometer interference in MEG as a spatially homogeneous magnetic field”. In: NeuroImage 244 (Dec. 2021), p. 118484. ISSN: 1053-8119. DOI: 10.1016/j.neuroimage.2021.118484. URL: https://www.sciencedirect.com/science/article/pii/S1053811921007576 (visited on 04/06/2026).

[40] Tim M. Tierney et al. “Optically pumped magnetometers: From quantum origins to multi-channel magnetoencephalography”. eng. In: NeuroImage 199 (Oct. 2019), pp. 598–608. ISSN: 1095-9572. DOI: 10.1016/j.neuroimage.2019.05.063.

[41] M. A. Uusitalo and R. J. Ilmoniemi. “Signal-space projection method for separating MEG or EEG into components”. en. In: Medical and Biological Engineering and Computing 35.2 (Mar. 1997), pp. 135–140. ISSN: 1741-0444. DOI: 10.1007/BF02534144. URL: https://doi.org/10.1007/BF02534144 (visited on 04/06/2026).

[42] Fulong Wang et al. “Exploring the Potential of SSVER-BCI Based on Contactless Measurement Using Optically Pumped Magnetometers”. In: IEEE Journal of Biomedical and Health Informatics (2025), pp. 1–12. ISSN: 2168-2208. DOI: 10.1109/JBHI.2025.3644887. URL: https://ieeexplore.ieee.org/document/11301600 (visited on 03/12/2026).

[43] Ruonan Wang et al. “Noise and artifact suppression in SQUID and wearable OPM-MEG: A systematic review of background, physiological, and Technical interference”. In: NeuroImage 318 (Sept. 2025), p. 121403. ISSN: 1053-8119. DOI: 10.1016/j.neuroimage.2025.121403. URL: https://www.sciencedirect.com/science/article/pii/S1053811925004069 (visited on 08/14/2025).

[44] Norbert Wiener. Extrapolation, Interpolation, and Smoothing of Stationary Time Series: With Engineering Applications. en. URL: https://direct.mit.edu/books/oa-monograph/4361/Extrapolation-Interpolation-and-Smoothing-of (visited on 07/24/2025).

